# Ecosystem service gradients at protected area borders reveal multiple patterns and prevalent management conflicts

**DOI:** 10.64898/2026.06.30.735453

**Authors:** Alberto González-García, Margot Neyret, Adrián López-Tejedor, Marie Caroline Prima, Sara Si-Moussi, Julien Renaud, Maya Gueguen, Sandra Lavorel

## Abstract

Protected areas cannot halt biodiversity loss in isolation; integrating them with surrounding human-dominated landscapes is critical. However, this integration is challenged by substantial landscape heterogeneity at their borders, hindering our understanding of cross-border changes in ecosystem service provision. We introduce a novel framework for characterizing these dynamics by analyzing ecosystem service gradients along protected area borders. For 16 protected areas in the French Alps, we assessed 12 ecosystem services using a mix of established biophysical models and novel connectivity-based models for mobile species. These were aggregated into three stakeholder-driven domains reflecting respectively rural, cultural, and urban management priorities. Automated polynomial regression analysis classified borders into five gradient types. The most common were ‘Decreasing Gradients’, representing a decline in ecosystem services outside the protected area, and ‘Increasing Gradients’, with the opposite pattern. Our analysis reveals these patterns are driven by specific landscape configurations, uncovering frequent trade-offs between the three management priorities, where, for instance, landscapes supporting rural priorities often degrade cultural and urban ones. We also identify key opportunities for synergies, by identifying areas where ecosystem services for all three priority domains increase simultaneously outside the protected area. This spatially explicit typology provides a powerful diagnostic tool for designing targeted interventions, such as prioritizing habitat restoration where ecosystem services decline or managing agricultural landscapes to mitigate conflicts across management priorities, supporting a more effective integration of protected areas into the wider landscape.

**Article impact statement:** Determining ecosystem service gradients at protected area borders reveals management trade-offs, guiding targeted spatial planning.

## 1. Introduction

Protected areas are the cornerstone of global biodiversity conservation (Geldmann et al., 2019; Gray et al., 2016). However, despite their successes, global biodiversity continues to decline at an alarming rate (Díaz et al., 2019; IPBES, 2019), emphasizing that a conservation strategy focused solely within protected area boundaries is insufficient. Even ambitious global targets, such as the Kunming-Montreal Framework’s goal to protect at least 30% of the planet, inherently underscore the critical importance of the vast landscapes outside protected areas (Obura, 2023). These human-modified landscapes, encompassing everything from managed forests and farmlands to urban centers, are essential for biodiversity conservation, landscape connectivity, and human well-being (Kremen, 2015; Kremen & Merenlender, 2018). Consequently, conservation paradigms are shifting towards integrated approaches that emphasize these crucial linkages (Cumming et al., 2015; Díaz et al., 2020), and global frameworks now explicitly call for the integration of sectors like agriculture and urban development into broader conservation strategies.

The long-term viability of protected areas is therefore intrinsically linked to the surrounding landscape, as they are not isolated islands but complex social-ecological systems (Cumming et al., 2015; Cumming & Allen, 2017). The effectiveness of the global protected area network depends fundamentally on its ability to support species movement through the surrounding matrix, which is essential for maintaining viable populations (Brennan et al., 2022). This landscape permeability also underpins the flow of ecosystem services to people, which can be diverse in nature: some are locally-determined stocks like carbon storage, while others are mediated by mobile organisms, like pollination (Martínez-Salinas et al., 2022), or conditioned by access like outdoor recreation (González-García et al., 2020). However, as spatial planning instruments, protected areas impose restrictions that generate unwanted dynamics at their boundaries, such as displacing land-use pressures (leakage) or attracting new urbanization (Ewers & Rodrigues, 2008; González-García et al., 2022). While landscape-scale principles suggest that securing these benefits requires maintaining significant semi-natural habitat in the matrix (Mohamed et al., 2024) our understanding of the fine-scale ecological and ecosystem service patterns across protected area borders remains in its infancy.

Recognition of this critical knowledge gap about protected area borders acknowledges that they are not uniform lines but highly heterogeneous interfaces where conservation and human activities collide (González-García et al., 2022). While research on landscape fragmentation has provided crucial insights into its impacts on biodiversity (Fahrig et al., 2019), the explicit link between landscape connectivity and the provision of ecosystem services has often been overlooked (Mitchell et al., 2015). Because administrative borders rarely coincide with natural boundaries, many ecosystem services are inherently transboundary and rely on spatial subsidies provided by mobile organisms across the landscape (López-Hoffman et al., 2017). Securing these benefits requires quantifying border gradients, which represent the spatial changes in ecosystem service provision with increasing distance from a border (Huang et al., 2026). However, these secondary contributions to surrounding areas are often inconsistently described due to a lack of interdisciplinary methodological frameworks (Cumming et al., 2025). Furthermore, because stakeholders perceive and manage nature as interconnected bundles of qualities rather than isolated benefits (Klain et al., 2014), defining these bundles based on social priorities is essential to identify the spatial trade-offs that drive landscape conflicts (Plieninger et al., 2019; Zoderer et al., 2019). Since overlooking the diverse values associated with these benefit bundles can undermine conservation efforts (Gross et al., 2025), untangling these complex interactions through consistent socio-ecological frameworks is critical for developing conservation strategies that are effective in the face of the intertwined biodiversity and climate crises.

Here, we introduce a novel framework to address this gap by quantitatively characterizing spatial variation of ecosystem services across protected area borders. By shifting the scale of analysis from the entire protected area to numerous individual border segments, our approach captures the high spatial variability of these interfaces, significantly advancing the state of the art in boundary research. Our framework aims to answer two central questions: (1) Do ecosystem service gradients at protected area borders form distinct, recurring patterns? And (2) What are the underlying landscape configurations that drive these patterns, and the resulting synergies and trade-offs between different stakeholder management priorities? We hypothesize that the contrast between conservation inside and human activities outside protected areas will frequently generate multiple patterns across ecosystem service bundles, resulting in management trade-offs. Furthermore, taking a spatially explicit approach, we expect these trade-offs to vary along the borders, directly driven by the specific landscape configurations of the surrounding matrix.

To answer these questions, we applied our approach to a network of protected areas in the French Alps, a region characterized by steep altitudinal gradients and a highly heterogeneous landscape matrix where protected areas are embedded within a complex mosaic of urbanized valleys, rural agricultural lands, and high mountain peaks. To capture this complexity, we advanced beyond standard assessments by using established biophysical models for services like carbon sequestration and flood regulation, alongside novel connectivity-based models for mobile, service-providing species such as pollinators and seed vertebrate dispersers. These ecosystem services were aggregated into three bundles reflecting rural, cultural, and urban management priorities, rooted in the region’s specific socio-ecological context (Vannier et al., 2019). A key innovation of our framework is the use of structural buffers that account for landscape permeability by integrating topography and land use. Unlike traditional Euclidean distances, this approach provides an ecologically realistic characterization of how landscape configuration mediates ecosystem service provision, allowing us to objectively classify border profiles into a typology of patterns.

## 2. Methods

We applied our framework to a network of 16 protected areas encompassing National Parks, Regional Nature Parks, and Nature Reserves across the French Alps (Fig. 1A; see Appendix S1 for detailed site characteristics). This region presents a landscape matrix dominated by forest cover and open alpine habitats at higher elevations, which transition into agricultural mosaics of pastures and crops on the slopes and valley floors, alongside expanding urban centers in the lowlands. To characterize the interfaces between surrounding land uses and protected zones, we followed a sequential workflow. First, we modeled 12 ecosystem services using two parallel approaches: established biophysical models and novel connectivity-based models for mobile species (Fig. 1B). These ecosystem services were then aggregated into three bundles reflecting distinct stakeholder-driven management priorities (rural, cultural, and urban). Subsequently, we analyzed the relative gradients for each bundle along border segments using structural buffers, which account for landscape permeability and topography to sample values from the interior to the exterior (Fig. 1C). Finally, we applied an automated classification procedure based on polynomial regression, which objectively identified the best-fitting model for each profile to define a typology of five gradient patterns (Fig. 1D).

**Figure 1.**
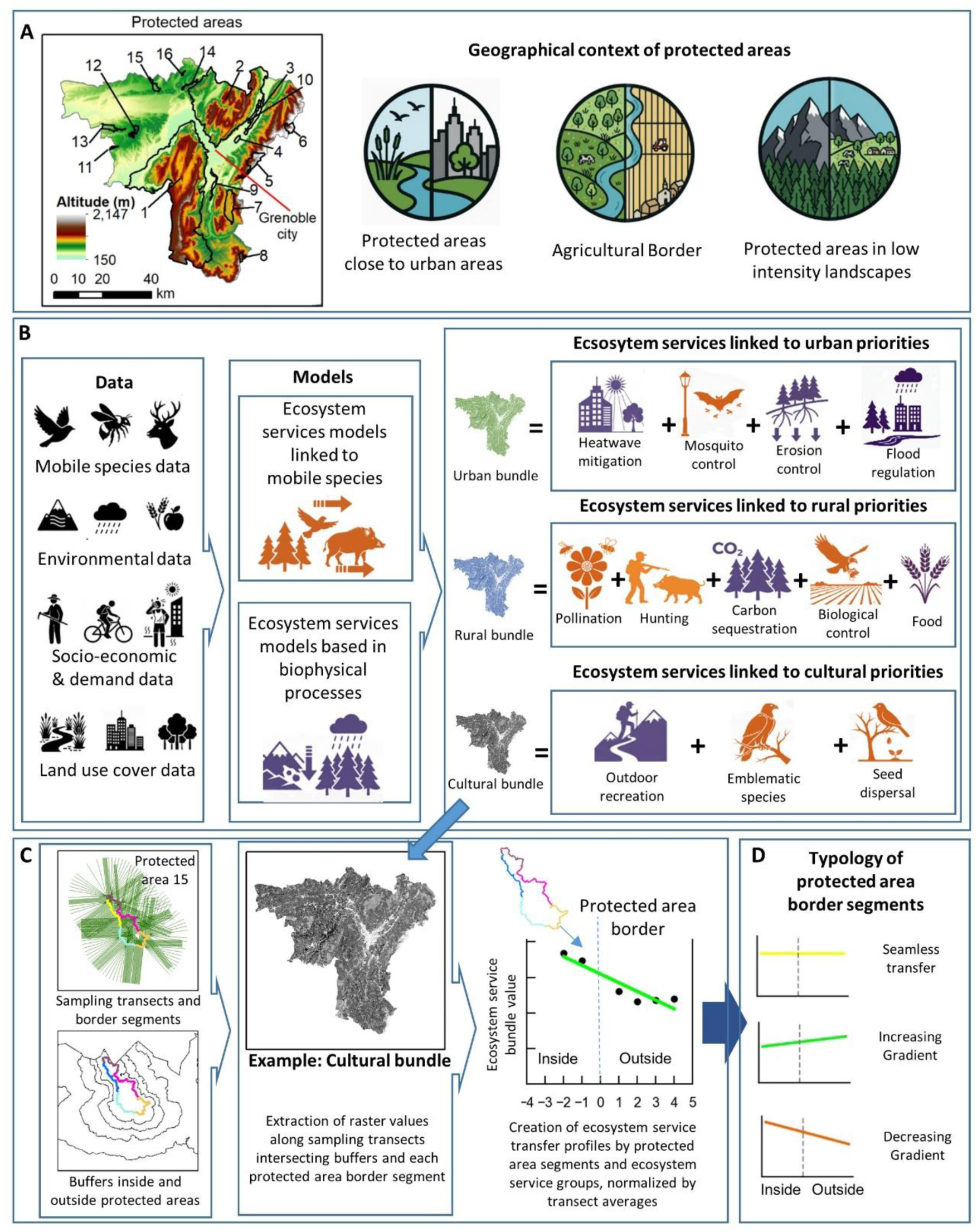
Methodological framework for characterizing ecosystem service gradients across protected area borders. (A) Our framework is applied to a diverse network of 16 protected areas in the French Alps, where borders are complex interfaces, illustrated here with three examples. (B) We first model 12 key ecosystem services by combining biophysical and connectivity-based models and integrating human demand criteria. These ecosystem services are then aggregated into three bundles reflecting distinct rural, cultural, and urban management priorities. (C) To analyze and compare the patterns of these ecosystem service bundles, we divide protected area boundaries into segments and sample the values of each of the three bundles along perpendicular transects. The analysis is structured using structural buffers (concentric zones of similar landscape permeability based on topography and land use) which serve as discrete steps for sampling, allowing us to construct a detailed gradient profile for each ecosystem service bundle at each segment. (D) Finally, these profiles are classified into a typology of five patterns, shown here with three illustrative examples. This comparative analysis of gradients across bundles allows us to identify and explain the synergies and trade-offs that are central to managing these complex interfaces.

### 2.1. Modelling ecosystem services

We modeled 12 ecosystem services selected for their critical relevance to the socio-ecological context of the French Alps, informed by prior regional stakeholder assessments (Vannier et al., 2019). The selection includes ecosystem services essential for mitigating natural hazards and health risks (erosion control, flood regulation, heatwave mitigation, mosquito control), ecosystem services central to the local economy (food production, supported by pollination and biological control), and ecosystem services linked to the region’s rich biodiversity and recreational pressure (outdoor recreation, hunting value, emblematic species, seed dispersal), along with carbon sequestration for its global relevance. To ensure spatial consistency across all ecosystem service models, we utilized a high-resolution (5m) land use and land cover map derived from Marsoner et al. (2023). We quantified these ecosystem services using two complementary modeling approaches: biophysical models and connectivity-based models for mobile species. Crucially, we incorporated spatially explicit human demand criteria where relevant to ensure the assessment captured the actual benefit to beneficiaries. For locally consumed ecosystem services like biological control or pollination, we restricted the analysis to areas where the ecosystem service is needed, such as crops or urban green spaces. For ecosystem services with global or diffuse beneficiaries, such as carbon sequestration, emblematic species, or seed dispersal, demand was considered ubiquitous across the landscape (see Table 1 for details in ecosystem services and demand rationale).

**Table 1.** Overview of the 12 ecosystem services modeled, specifying the methodological approach, the indicators used to quantify supply, and the spatial criteria applied to integrate human demand and access.

| Ecosystem service | Modeling approach | Indicator of supply | Integration of demand |
| --- | --- | --- | --- |
| Food production | Biophysical (yield statistics) | Energy equivalent of crop production (GJ/ha) | Supply restricted to productive agricultural parcels. |
| Carbon storage and sequestration | Biophysical (InVEST) | Carbon stored in biomass and soil (Mg C/ha) | Global climate regulation ecosystem service; demand considered ubiquitous. |
| Heatwave mitigation |  | Cooling capacity index | Heat island effect intensity in populated areas. |
| Flood regulation |  | Runoff retention index | Reference precipitation events requiring retention in urban catchments. |
| Erosion control |  | Sediment retention | Sediment export to streams affecting water quality downstream. |
| Outdoor recreation | Biophysical (Regional model) | Landscape attractiveness index | Accessibility and visitation density (crowd-sourced GPS tracks). |
| Pollination | Connectivity-based (Omniscape) | Pollinator potential connectivity | Intersection with pollination-dependent crop areas. |
| Biological control |  | Predator potential connectivity (78 species) | Intersection with crop areas and urban green spaces. |
| Mosquito control |  | Predator potential connectivity (81 species) | Intersection with human settlements and mosquito presence records (*Aedes albopictus*, *Culex pipiens*). |
| Hunting value |  | Game species potential connectivity (24 species) | Accessibility regulations (excluding reserves and proximity to buildings). |
| Seed dispersal |  | Frugivorous potential connectivity (81 species) | Beneficial for overall ecosystem function; no specific spatial restriction. |
| Emblematic species |  | Iconic species potential connectivity (96 species) | Cultural existence value; beneficial across the entire landscape. |

#### 2.1.1. Biophysical ecosystem service models

We used established biophysical approaches for six ecosystem services. Food production was quantified at the pixel level based on regional crop yield statistics converted to energy equivalents following Lasseur et al. (2018). Outdoor recreation was assessed using a regional model that incorporates landscape attractiveness, accessibility, and crowd-sourced GPS track data to represent realized recreational use (Byczek et al., 2018). The remaining four ecosystem services were modeled using the InVEST software suite (Natural Capital Project, 2023). Carbon storage and sequestration was calculated by summing carbon stocks in four pools (aboveground biomass, belowground biomass, soil, and dead organic matter) for each land cover type. Heatwave mitigation was estimated using the Urban Cooling model, which calculates a cooling capacity index based on shade, evapotranspiration, and albedo. Flood regulation was assessed via the Urban Flood Risk Mitigation model, which estimates runoff retention potential based on the Curve Number method derived from soil and land use characteristics. Erosion control was modeled using the Sediment Delivery Ratio model, which computes sediment retention based on topography, rainfall erosivity, and soil erodibility following the Revised Universal Soil Loss Equation. Specific parameters, data sources, and demand integration details for each of these six models are provided in Appendix S2.

#### 2.1.2. Connectivity-based ecosystem service models

For ecosystem services provided by mobile organisms, specifically pollination, hunting value, biological control, seed dispersal, mosquito control, and emblematic species, we developed a multi-step modeling approach that explicitly incorporates species movement across the landscape matrix. First, we developed species-specific habitat suitability models at a high resolution (5m) for 224 vertebrate species using an ensemble of machine-learning algorithms (Si-Moussi & Thuiller, 2024; O’Connor et al., 2024) and integrating species-specific habitat preferences (Maiorano et al., 2020). Second, we modeled functional landscape connectivity for each species using the Omniscape algorithm (McRae et al., 2016). To do so, we transformed habitat suitability into landscape resistance surfaces using a non-linear function (Keeley et al., 2016) and applied circuit theory to identify source habitats and calculate current flow (Prima et al., 2024). Finally, ecosystem service maps were generated by aggregating the connectivity-corrected habitat suitability maps of all species contributing to a given function. To map the actual ecosystem service provision, we intersected these connectivity maps with spatial demand layers where appropriate. For example, the pollination ecosystem service was restricted to agricultural areas dependent on pollinators, while hunting value was spatially refined by excluding areas where hunting is prohibited, such as strict reserves and safety buffers around human settlements. The rationale for species grouping and detailed modeling parameters are provided in Appendix S3.

### 2.2. Aggregation into management priorities ecosystem service bundles

To analyze patterns of synergies and trade-offs, we grouped the 12 normalized ecosystem service maps into three bundles. This aggregation was not derived from statistical clustering of the biophysical maps (Mouchet et al., 2015), but was grounded in stakeholder priorities identified through long-term transdisciplinary research in the region (Lavorel et al., 2016; Crouzat et al., 2016; Vannier et al., 2019). These bundles, which cut across multiple values of nature for people (Pascual et al. 2017), represent how local stakeholders, including farmers, foresters, urban planners, and conservationists, perceive the interactions between ecosystem services and their management priorities. Defining bundles based on these priorities is a robust approach to uncover spatial trade-offs and synergies that purely biophysical models often underestimate (Plieninger et al., 2019). This choice reflects the evidence that stakeholders rarely perceive ecosystem services as isolated benefits, but rather as interconnected ‘bundled qualities’ (Klain et al., 2014). Recent empirical research in complex social-ecological systems confirms that stakeholder preferences naturally cluster into distinct domains, justifying the use of these associated benefit sets as a basis for conservation planning (Gross et al., 2025). By grouping services into these priority domains, our framework provides a diagnostic tool to identify spatial mismatches between conservation goals and human needs, which are at the root of most landscape conflicts at protected area borders (Zoderer et al., 2019).

The Rural bundle reflects priorities linked to agricultural production and rural livelihoods. It includes ecosystem services that support the farming economy (food production, biological control, pollination) and traditional practices (hunting value), as well as carbon sequestration, which stakeholders associate with the management of forests and the extensive rural matrix. The Urban bundle comprises ecosystem services primarily demanded by denser human populations to regulate natural hazards affecting infrastructure and health. This includes flood regulation, erosion control, and recent concerns for heatwave mitigation and mosquito control. The Cultural bundle aggregates ecosystem services contributing to recreational uses, heritage, and landscape identity across the region and its various ecosystems. It includes outdoor recreation and emblematic species, which underpin the region’s tourism economy and cultural attachment to nature. We also included seed dispersal in this bundle, as it is perceived as a foundational process maintaining the long-term integrity of the high-biodiversity landscapes that support these cultural values. Each bundle map was created by calculating the unweighted mean of its constituent normalized ecosystem service maps.

### 2.3. Ecosystem service gradient analysis and border typology

To analyze ecosystem service gradients, we developed structural buffers based on landscape resistance rather than Euclidean distance. This approach accounts for how topography and land cover facilitate or impede movement, capturing the functional connectivity across the border. We generated a resistance surface combining slope and land-use resistance values (detailed in Appendix S4.1). Using a cost-distance algorithm, we delineated five concentric zones of increasing accumulated resistance extending both inwards and outwards from the protected area boundaries (Fig. 1c), creating a consistent spatial framework to sample gradients across heterogeneous landscapes.

To characterize the interface, we divided the borders of the 16 protected areas into segments (ranging from 1.2 to 11.2 km in length) to capture variations along the perimeter. Within each segment, we generated perpendicular transects at 25 m intervals to sample ecosystem service bundle values across the structural buffers. To ensure spatial validity, we applied an automated filtering process (see Appendix S4.2, Figure S4 for specific parameters). This included truncating transects where they intersected with the buffer zones of neighboring protected areas to prevent the gradient from reflecting patterns linked to proximate protected areas, as well as discarding lines that were too short or topologically invalid. Finally, we normalized the values relative to each transect mean, allowing us to characterize the relative variation of ecosystem services across the buffer zones compared to the average value of the entire transect.

We constructed a representative gradient profile for each segment and bundle by averaging values from valid transects within each structural buffer zone (Fig. 1D). To classify these profiles into the A-E typology, we developed an objective procedure based on polynomial regression (Appendix S4.3). We fitted both first-degree (linear) and second-degree (quadratic) models to each profile, selecting the best-fit based on the minimum Mean Squared Error (MSE). We then applied specific thresholds to the model coefficients to distinguish between patterns: linear trends (Types A, B, and C) were defined by an MSE < 0.05 and classified based on their slope (flatness threshold < 0.01), while quadratic models (Types D and E) were selected to capture non-linear border effects—such as interface peaks or depressions—based on the sign of the quadratic term. This algorithmic approach ensures a replicable analysis that avoids the subjectivity of visual classification.

To identify the landscape drivers of these patterns, we analyzed three key components of landscape configuration inside and outside the protected area for each segment: composition (proportions of natural, semi-natural, and artificial land cover), structure (Shannon diversity index), and configuration (Moran’s I as a proxy for fragmentation). We used Linear Mixed Models (LMMs) to test whether changes in these metrics across the border explained the observed ecosystem service patterns (Appendix S5). In these models, we evaluated the significance of the interaction between border location (inside vs. outside) and gradient type, accounting for the non-independence of spatial data by including Protected Area and Border Segment as nested random effects.

### 2.4. Robustness analysis

To ensure that the aggregated management bundles accurately reflect the underlying patterns of individual ecosystem services, we conducted a three-level robustness analysis across all border segments. First, we evaluated geometric similarity by calculating the Pearson correlation (r) between the reconstructed curves of each individual ecosystem service and its corresponding bundle. Second, we assessed categorical agreement by identifying matches between the gradient typology labels (A–E) assigned at the ecosystem service level versus the bundle level. Finally, we performed a component influence analysis by re-aggregating an alternative version of the rural bundle that excluded place-based ecosystem services (food production and carbon storage). This third level allowed us to analyze the shift of border segments between typology categories and evaluate the specific weight of connectivity-based ecosystem services in driving the final classification patterns. These complementary evaluations provide a robust validation of the framework’s representativeness and diagnostic power (detailed methodology in Appendix S6).

## 3. Results

### 3.1. Typology of protected area borders

The classification of the 312 segment–bundle profiles (the 104 border segments analysed across the three management bundles) based on their relative patterns of ecosystem service change resulted in five distinct gradient types (Fig. 2). Overall, the ‘Decreasing Gradient’ (Type C) was the most frequent pattern, accounting for 49% of all profiles, followed by the ‘Increasing Gradient’ (Type B, 25%) and the ‘Flat Gradient’ (Type A, 9%). These linear patterns collectively describe over 80% of profiles. The remaining profiles were classified into two more complex, non-linear types: ‘Boundary Depression’ (Type D, 11%) and ‘Interface Peak’ (Type E, 5%). See Appendix S7.1 for further details (Figure S5 for border segments classification example). A consistency analysis confirmed the robustness of this classification, showing that the aggregated bundle-level patterns are representative indicators of component individual ecosystem service gradients. While local divergence occurs in the rural bundle, the assigned typology captures the dominant spatial trend across 97.7% of all bundle observations, providing a reliable basis for management diagnosis (Appendix S7.2, Table S9 and Figure S10). This validation shows that while the cultural bundle exhibits high internal consistency (median r = 1.00), the rural bundle is more heterogeneous, with specific ecosystem services like food production often diverging from the overall bundle trend. Nevertheless, the assigned typologies accurately reflect the dominant spatial trends across 97.7% of the analyzed cases, confirming the bundles are robust indicators for identifying general management patterns along the borders.

**Figure 2.**
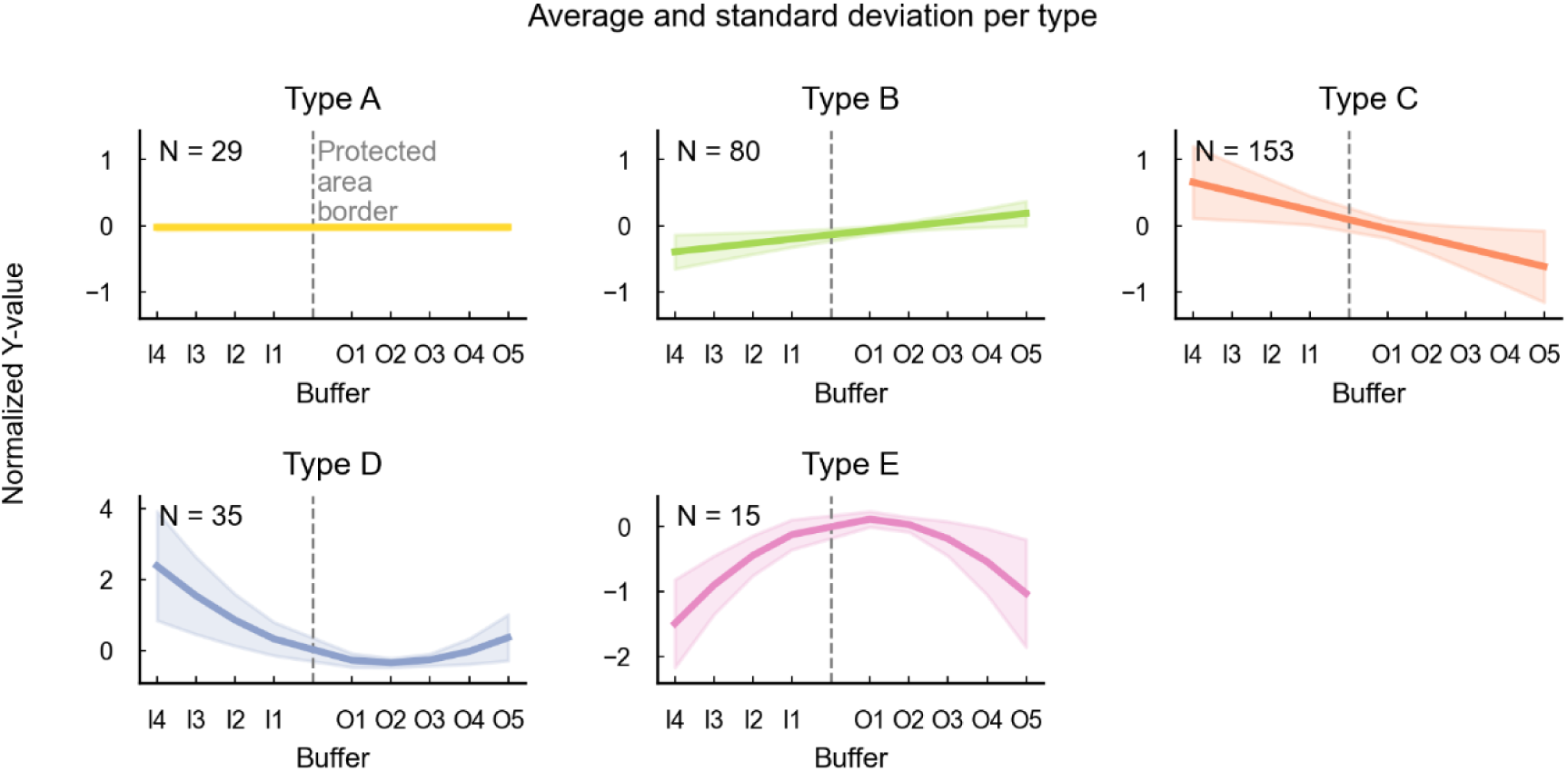
Typology of protected area borders based on ecosystem service bundle profiles. The profiles illustrate the normalized ecosystem service bundle values (Y-axis) across structural buffer zones from inside to outside protected areas (X-axis). Y-axis values represent the average ecosystem service bundle value within each buffer zone, normalized by the average value along the entire border segment transect line. This normalization highlights the relative ecosystem service bundle value within each buffer zone compared to the overall transect average value. The X-axis labels use “I” before the number to indicate zones inside the protected area and “O” before the number to indicate zones outside the protected area. The number of segment–bundle profiles classified into each typology (N) is indicated within each panel. The total number of border segments is 104, but since profiles are shown for three analyzed ecosystem service bundles (rural, cultural, and urban), the figure presents a total of 312 segment–bundle profiles. Figure S3 in the Supporting Information shows an example of the polynomial regressions generated for all the bundles and protected area border segments, including statistically significant values and detailed mean square errors.

Beyond their statistical classification, these five types represent distinct patterns of relative ecosystem service provision across the border (Fig. 2). Type A shows values consistently close to the transect mean, indicating a seamless transition. Type B reveals a positive trend, where ecosystem service provision becomes gradually higher than the transect mean outside the protected area. Type C, the dominant pattern, shows a negative trend, with ecosystem service provision consistently dropping below the transect mean outside the protected area. Types D and E represent more complex patterns, highlighting specific zones of significant deviation from the mean, such as depressions or peaks right at the border. The following sections analyze the landscape configurations that drive this typology (Fig. 3) and how the three bundles combine along borders to produce synergies and trade-offs (Fig. 4).

**Figure 3.**
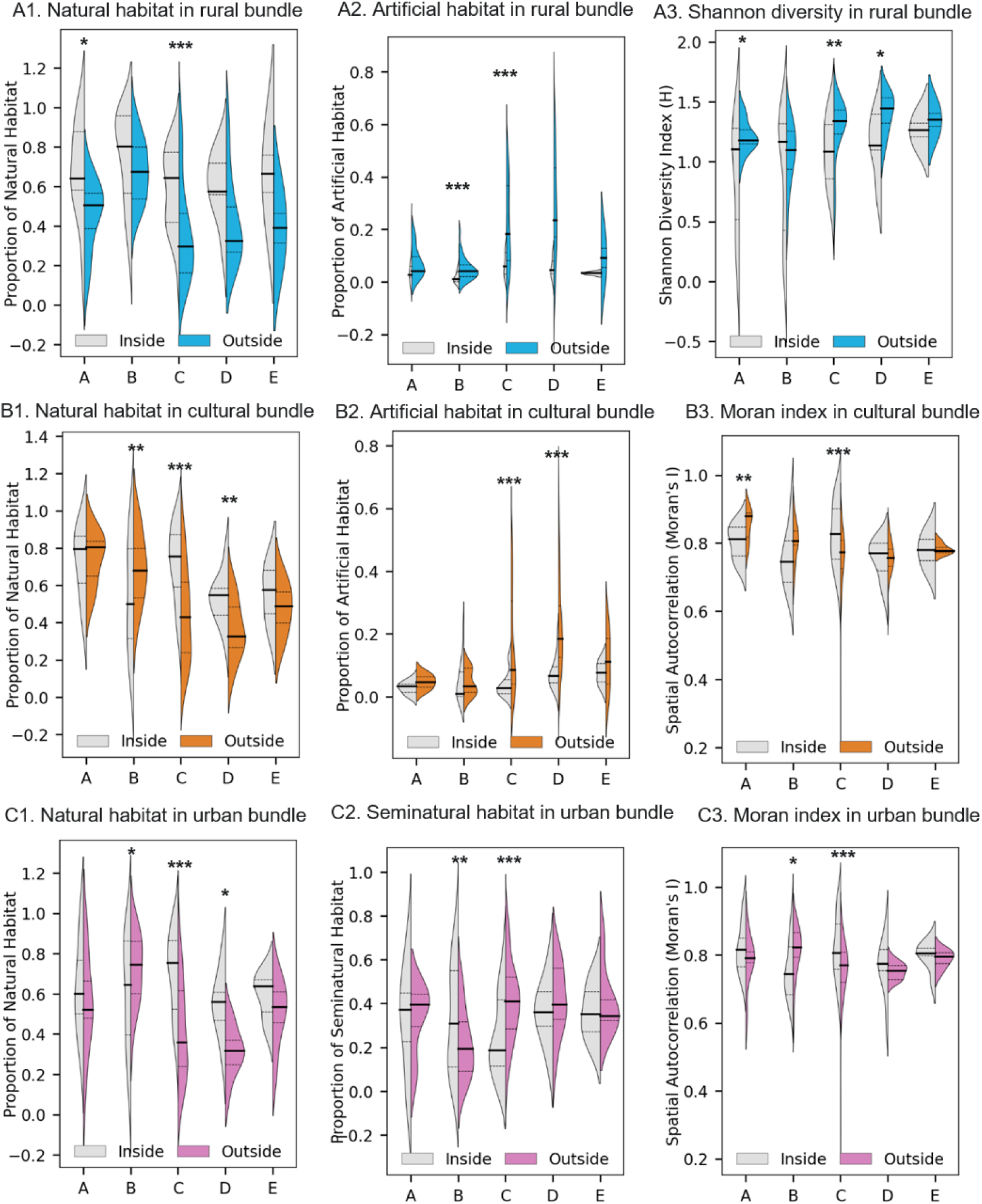
Landscape signature of each border type, shown separately for the three ecosystem service bundles. For every border type (A–E) defined in Figure 2, this figure characterises the landscape composition and configuration on each side of the protected area boundary, revealing the landscape conditions that give rise to each gradient type. The figure is organised in three rows, one per bundle: (A) Rural, (B) Cultural, (C) Urban. In each panel, the y-axis is the landscape metric named in the panel title, the proportion of natural, artificial or semi-natural habitat, the Shannon diversity index (H), or spatial autocorrelation (Moran’s I, a proxy for fragmentation, where higher values indicate a more continuous, less fragmented landscape). Violin plots compare the distribution of that metric for areas inside (grey) versus outside (coloured) the protected area, with border types (as classified for the corresponding bundle) on the x-axis; because each segment is classified independently per bundle, a given type need not comprise the same segments across rows. Read alongside Figure 2, the figure shows the landscape basis of each gradient: for example, the outward decline in ecosystem service provision of Type C coincides with more natural habitat inside the boundary and more artificial habitat outside it. Solid and dashed lines within violins represent the median and interquartile range, respectively. Asterisks indicate a statistically significant difference between the inside and outside distributions for a given border type (Wilcoxon signed-rank test; * p < 0.05, ** p < 0.01, *** p < 0.001).

**Figure 4.**
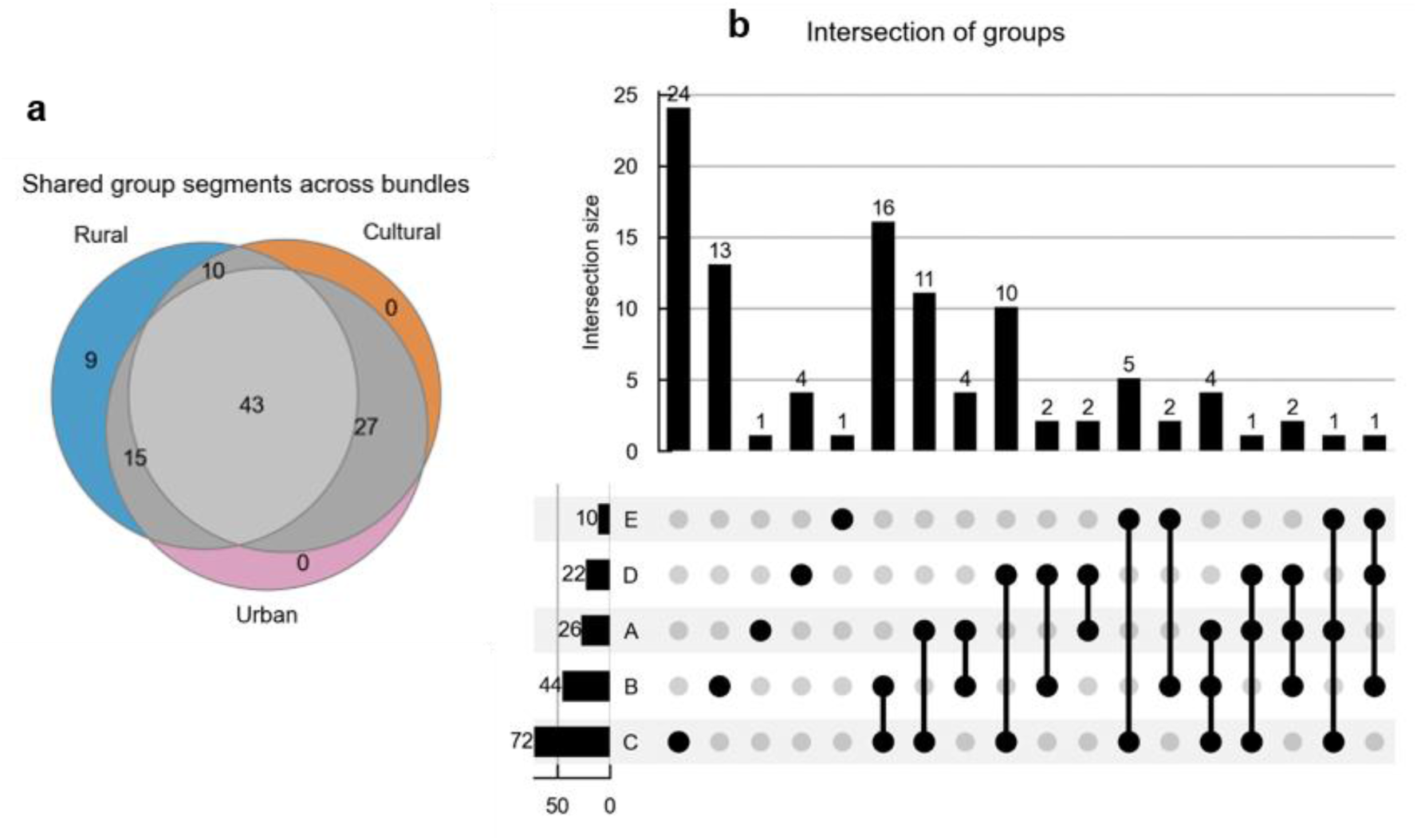
Agreement and divergence of border types across the three ecosystem service bundles. For each of the 104 border segments, the type (A–E) assigned to the rural, cultural and urban bundles is compared. (a) Venn-style diagram summarizing how often segments share the same type across bundles; each segment is counted once, so the values sum to 104. The central value (43) is the number of segments classified as the same type across all three bundles simultaneously (e.g., all three Type A, or all three Type B). (b) UpSet plot of the most frequent combinations of types across the 104 segments. Vertical bars (intersection size) give the number of segments sharing a specific combination of types across their three bundles; each segment contributes to exactly one combination, so these bars also sum to 104 (e.g., the leftmost bar: 24 segments are Type C in all three bundles). Horizontal bars (set size) give, for each type, the number of segments in which that type appears in at least one of the three bundles; because a single segment can carry different types across its bundles, it is counted in every set to which it belongs, and the set sizes therefore sum to more than 104. Each set size counts segments, whereas the per-type totals in Figure 2 count the 312 segment–bundle profiles; the two use different units, which is why their values differ. Matrix of dots: a filled dot indicates that the type in that row is part of the combination in the column above; connecting lines group the types forming each combination.

### 3.2. Landscape drivers of border patterns

To understand the mechanisms underpinning our typology, we analyzed landscape composition and configuration across the border for each bundle using Linear Mixed Models. The results reveal that each border type corresponds to a distinct landscape signature, explaining the observed ecosystem service patterns (Fig. 3). See Appendix S7.1, Figure S6 and S7 for detailed ecosystem services models and Figure S8 for an example of ecosystem services distribution around protected areas. See Appendix S7.3, Table S10 for a full statistical summary of these results.

For the rural bundle, patterns are primarily driven by the transition between natural and agricultural landscapes, with urban encroachment acting as a critical factor. Type A (’Flat Gradient’) borders are characterized by a significant decrease in natural habitat outside the protected area (p < 0.05) compensated by an increase in semi-natural and agricultural land, while landscape diversity increases (Fig. 3A1, 3A3). Type B (’Increasing Gradient’) shows relative landscape homogeneity across borders, suggesting low-intensity farming mixed with natural elements. The most pronounced change occurs for Type C (’Decreasing Gradient’) borders, where natural habitat declines sharply (p < 0.001) and is replaced by significantly increasing artificial land (p < 0.001) (Fig. 3A1, 3A2). The clarity of these linear patterns is strongly influenced by two locally-determined ecosystem services: food production and carbon storage. The component influence analysis (Appendix S7.4 and Table S11) confirms that excluding these two services shifts the border profiles from simple linear gradients toward more complex, non-linear patterns. This result reinforces that food production and carbon storage act as the dominant spatial signals for the rural bundle, defining the broad trends within which non-linear patterns of connectivity-based ecosystem services are embedded.

For the cultural bundle, ecosystem service provision is linked to landscape integrity. The decreasing gradient of Type C is explained by the replacement of natural habitat by both agricultural and artificial land (Fig. 3B1, 3B2). Such landscapes are less suitable for emblematic species or seed dispersal. This landscape change results in a more uniform matrix outside the protected area, with a significant decrease in fragmentation (Fig. 3B3, p < 0.001).

The patterns for the urban bundle reveal that high ecosystem service provision is associated with managed mosaics rather than just landscapes with high natural cover. The ‘Increasing Gradient’ (Type B) corresponds with a landscape outside the protected area that is more natural (p < 0.05), less semi-natural (p < 0.01) and more fragmented (p < 0.05) (Fig. 3C1, 3C3). This points to a ‘managed natural mosaic’, such as pastures with scattered trees, that favors regulating ecosystem services. Conversely, the decreasing gradient (Type C) is primarily explained by the substitution of natural habitat with agricultural land (p < 0.05 for both, Fig. 3C1, 3C2). This indicates that the expansion of agriculture alone is sufficient to degrade key biophysical functions, leading to a more spatially continuous landscape (Fig. 3C3, p < 0.001).

### 3.3. Cross-bundle synergies and trade-offs in border patterns

The distinct landscape drivers for each bundle lead to varying combinations across borders, resulting in both synergies and trade-offs for management. While a large number of segments (43) share a common border type across all three bundles, many others present divergent patterns, highlighting the prevalence of potential management conflicts (Fig. 4a). A detailed analysis of the most frequent combinations (Fig. 4b) reveals that the most common pattern is a consistent ‘Decreasing Gradient’ (Type C) across all three bundles (24 segments). This shared decline, typically found in landscapes dominated by intensive agriculture or urban expansion, suggests that targeted restoration could yield “triple-win” outcomes. For example, diversifying homogeneous agricultural or urban areas with green infrastructure like hedgerows and parks could simultaneously enhance regulating, cultural, and even some rural ecosystem services.

Synergies are also apparent in the 13 segments showing a consistent ‘Increasing Gradient’ (Type B) across all domains. However, the most frequent conflict (16 segments) is the trade-off between an Increasing Gradient for the rural bundle and a Decreasing Gradient for the urban and cultural bundles. Our robustness analysis (Appendix S7.2) quantifies the mechanics of this conflict: while the cultural bundle behaves as a homogenous geometric signal (median r = 1.00), the rural bundle is a composite of diverging patterns. Specifically, the provision of food production and biological control shows near-zero correlation with the overall bundle trend (median r = −0.06 and 0.03, respectively), indicating they follow spatial signatures that are independent of regulating and cultural functions. This confirms that these production-oriented services increase in specific landscape configurations where urban and cultural benefits simultaneously decline, highlighting a structural socio-ecological trade-off. Such patterns point toward opportunities for nuanced management where habitat patches could be integrated into agricultural production land to maintain the rural bundle, while also enhancing regulating functions for urban dwellers. The spatial distribution of these key combinations is detailed in Figure S9 and S11 Appendix S7.

## 4. Discussion

### 4.1. From landscape signatures to general principles for management

The management of protected area borders exemplifies a “wicked problem,” characterized by complexity and conflicting priorities among stakeholders (DeFries & Nagendra, 2017). Our results confirm this: most border segments present divergent patterns across bundles, highlighting that management conflicts are more prevalent than synergies. Our framework provides a spatially-explicit map for detecting these conflicts that reflect different stakeholder priorities, offering a foundation for moving towards more nuanced, context-specific management.

Our analysis reveals a prevalent trade-off between the rural bundle and the urban and cultural bundles, a finding that reflects the challenge of creating landscapes that work for biodiversity and people (Kremen & Merenlender, 2018). The core conflict arises from differing land-use logics: an intensive production logic (agriculture) versus a strict conservation-based logic that relies on intact habitats (Robalino et al., 2017). Our results surpass this polarization by adding a crucial structural dimension to this trade-off. Landscapes supporting an ‘Increasing Gradient’ (Type B) for the rural bundle, which includes connectivity-based ecosystem services like pollination and biological control, are often characterized by small-scale mosaics of natural and semi-natural habitats. Our robustness analysis (Appendix S7.2) confirms that these specific services follow spatial patterns that are independent of the bundle’s average trend (r < 0.35), as their provision depends on landscape heterogeneity rather than the total amount of natural land cover. This aligns with evidence that habitat fragmentation *per se*, independent of habitat loss, can have neutral or even positive effects on certain taxa by increasing landscape heterogeneity (Fahrig et al., 2019). Our framework captures this complexity, showing how these fragmented configurations can support mobile species that provide ecosystem services critical to rural livelihoods, while simultaneously degrading the larger, more continuous habitats required by ecosystem services in the cultural bundle. Our findings for the cultural bundle indicate that the reduction in landscape fragmentation at certain borders reflects a shift toward large-scale, homogeneous land uses, such as intensive agriculture or expanding urban centers. These structural changes likely degrade the ecological conditions and aesthetic qualities necessary to support emblematic species and recreational value.

However, this conflict is not inevitable. The literature provides clear principles for managing these “working landscapes” to mitigate trade-offs. Adopting “fine-grained landscape” strategies that enhance configurational heterogeneity (e.g., smaller fields, higher edge density, hedgerows or agroforestry) is a key structural pathway to support multifunctionality (Hass et al., 2018; Lavorel et al., 2022). This approach can reverse the shared decline (Type C) we observe in homogeneous landscapes and transform conflictive borders. This aligns with emerging principles for area-based conservation (Riva et al., 2024), which increasingly focus on managing the matrix to mitigate the inevitable trade-offs between different ecosystem service bundles and biodiversity (Neyret et al., 2025). For instance, integrating green infrastructure could sustain rural livelihoods while enhancing regulating and cultural ecosystem services (Schulte et al., 2017).

Despite the prevalence of trade-offs, our results also identify “win-win” synergies. This aligns with global evidence that positive conservation and socioeconomic outcomes are often synergistic (Oldekop et al., 2016). We propose that our methodology can pinpoint these bright spots (Frei et al., 2018), identifying them as ‘living labs’ to understand the underlying contextual factors, such as local governance structures or historical land-use legacies, that our models do not capture and that are crucial for replicating success (Daw et al., 2015).

### 4.2. A diagnostic framework for spatial planning and management priorities in socio-ecological interfaces

A primary innovation of our framework is the direct integration of ecosystem services demand. While many analyses focus on ecosystem service biophysical supply, management priorities arise from the interaction between that supply and human needs (King et al., 2015). This approach aligns with recent ‘dual-zone’ governance models (Li et al., 2026), which demonstrate that linking human functional zones with ecosystem service bundles is essential for the precise identification of sensitive areas. Our results validate this necessity by showing that management priorities cannot be effectively addressed without understanding how human needs mediate the biophysical supply at the border. By incorporating demand indicators, such as the location of specific crops for pollination or proximity to settlements for mosquito control, our method captures the demand dimension of ecosystem services that is fundamental for management. This approach avoids potential mismatches between where ecosystem services are produced and where they are needed (González-García et al., 2020). For instance, a landscape connectivity map for mosquito predators might show high values in remote forests, but by weighting this by the co-occurrence of mosquitoes and people, our model identifies critical areas for conservation at complex urban-rural interfaces. This allows managers to pinpoint where specific interventions, like reducing light pollution to favor bats, would be most effective.

This integrated approach enhances the diagnostic power of the framework, allowing it to deconstruct the drivers of bundle patterns. Following the call for more mechanistic approaches to bundle analysis (Dade et al., 2019; Meacham et al., 2022), our analysis of place-based ecosystem services demonstrates this capability (Appendix S7.4). It reveals that many apparently uniform ‘Flat Gradient’ borders (Type A) in the rural bundle are a “false equilibrium”, an artifact of the spatial compensation between high carbon storage inside the protected area and high food production outside. Without this deconstruction, a simple correlational analysis would conclude to stability and miss the underlying land-use trade-off (Crouzat et al., 2015). Furthermore, the framework’s diagnostic utility extends to the individual ecosystem service level. While our study focuses on the French Alps, the conceptual and methodological framework is adaptable to any socio-ecological context where protected area boundaries face land-use pressure. This remains relevant in regions with legally established buffer zones; in such cases, our gradient analysis provides a diagnostic tool to evaluate if these administrative zones are functionally effective or if ecosystem service provision drops abruptly regardless of the legal limit.

Ultimately, by revealing the complexity and prevalence of trade-offs informed by the combination of ecological processes and societal demand, our framework is well-suited to inform deliberative planning processes. The spatial identification of where different ecosystem service bundles come into conflict provides a tangible, evidence-based foundation for stakeholder negotiation (Shipley et al., 2020). As our analysis is based on bundles that aggregate ecosystem services according to stakeholder priorities (i.e., ‘rural’, ‘urban’, and ‘cultural’ domains), rather than on purely statistical correlations, it is particularly adept at uncovering the real-world trade-offs that are central to land-use planning (Martín-López et al., 2012; Vannier et al., 2019). For instance, a stakeholder may live in the city but prioritize ecosystem servicers associated with the cultural bundle in remote mountain landscapes. Our framework captures these nuances, allowing results to serve as a common reference point for stakeholders to discuss the consequences of different land-use scenarios (Longato et al., 2021) and to co-design more adaptive and socially robust management strategies (Rounsevell et al., 2021). For instance, in segments where we identified a ‘Boundary Depression’ (Type D), planners should first evaluate the underlying cause. Where these patterns reflect human-induced degradation, restoration should be prioritized to reconnect fragmented matrices. However, in cases where the depression is driven by inherent biophysical barriers, such as cliffs or steep rock faces, managers must instead account for these natural discontinuities to ensure that spatial planning remains grounded in the region’s physical constraints. Conversely, in agricultural corridors where the trade-off between rural and cultural bundles is sharpest, our typology suggests that implementing targeted agri-environmental schemes could enhance landscape heterogeneity, potentially transforming a ‘Decreasing Gradient’ (Type C) into a more synergistic transition that maintains both production and cultural heritage (Lomba et al., 2020; Mupepele et al., 2021).

### 4.3. Methodological advances, limitations, and future directions

While our framework introduces several methodological advances, we acknowledge its limitations. A full empirical validation of such a complex, multi-service framework is a considerable challenge. A critical consideration for replicating our approach lies in the definition of ecosystem service bundles. Our aggregation into rural, urban, and cultural bundles was grounded in extensive prior transdisciplinary research in the region (Lavorel et al., 2016; Crouzat et al., 2016; Vannier et al., 2019), which allowed us to identify distinct priority domains perceived by local stakeholders. Replicating this framework in other socio-ecological systems requires a similar foundational understanding of local priorities; applying our specific bundle composition elsewhere without this context could lead to misinterpretations (Martín-López et al., 2012). Furthermore, while we assigned each ecosystem service to a primary bundle to facilitate the analysis of trade-offs, we acknowledge that individual ecosystem services often cut across priority domains—for instance, the value of emblematic wildlife species is recognized by rural and urban stakeholders, but not specific to these domains (Lavorel et al., 2016).

Regarding the individual models, our approach to integrating demand is most effective for ecosystem services with local beneficiaries. For globally-driven ecosystem services like carbon storage and food production, whose demand dynamics are highly complex and delocalized, we adopted a necessary simplification by mapping their supply only (Martín-López et al., 2019). We also acknowledge specific constraints in the biophysical models (detailed in Appendix S8). For instance, our carbon stock model relies on data extrapolation from similar Alpine regions, and the heatwave mitigation model, while providing a valid biophysical index of cooling capacity, applies a tool originally designed for urban environments to the broader landscape. Additionally, our analysis focuses on patterns at the immediate border interface (within a 4 km transect extent). Consequently, patterns such as the ‘Increasing Gradient’ (Type B) should be interpreted as local trends at the interface scale, as these gradients would inevitably plateau or decline at broader distances from the protected area.

These limitations open avenues for future research, particularly in the context of global environmental change. We are currently using this diagnostic framework to inform participatory scenario planning with local stakeholders to explore how future land-use trajectories could alter the synergies and trade-offs (Rodríguez et al., 2023). This prospective approach is crucial because the border patterns we identified are not static, and the same landscape connectivity that supports benefits might also facilitate the spread of undesirable phenomena like wildfires or agricultural pests (Plowright et al., 2021; Sá et al., 2022). The resilience of biodiversity in the face of climate change will largely depend on the ability of species to move across these “working landscapes” (Kremen & Merenlender, 2018). Our findings confirm that many protected area borders, far from being permeable, are zones of intense land-use conflict that create barriers to this necessary movement (Haddad et al., 2015). Such tensions are expected to intensify as climate change forces species to shift their ranges beyond protected area borders, increasing the potential for human–wildlife conflicts along these borders (Abrahms et al., 2023). Therefore, a diagnostic framework like ours, which allows managers to identify, map, and understand the mechanisms of these border conflicts, becomes an essential foundation for proactive climate adaptation planning (Andersson et al., 2025).

## 5. Conclusions

This study introduces a novel framework for analyzing the interfaces of protected area borders. We move beyond broad-scale assessments to demonstrate that the provision of ecosystem services at these borders organizes into a predictable, fine-scale typology of gradients. Our key finding is not simply that trade-offs exist, but that they manifest as a complex spatial mosaic of conflict and synergy along the same border, representing a key management opportunity. We show that landscapes configured to support rural management priorities frequently degrade ecosystem services associated with the urban and cultural domains, but we also identify specific locations where synergies across all three domains occur. By linking these patterns to underlying landscape configurations, our framework acts as a powerful diagnostic tool. It provides the spatially explicit evidence needed to move beyond “one-size-fits-all” management and instead design targeted interventions that reconcile multiple objectives. This allows for the design of context-specific strategies, such as implementing green infrastructure where conflicts are sharpest, or studying the governance of synergistic areas to replicate success, that are essential for effectively integrating protected areas into multifunctional, sustainable landscapes in the face of global change.

## Supporting information

Appendix_Gonzalez_Garcia_et_al_2026

## Author Contributions

A.G.G, M.N, and S.L designed and performed the research. A.G.G, M.N, M.C.P, S.S, J.R, and M.G, created and prepared the data and models. A.G.G and A.L.T, developed the code and methods. A.G.G, A.L.T and S.L analyzed the data. A.G.G and S.L wrote the paper.

## Acknowledgments

Project RECONNECT, Biodiversa+, the European Biodiversity Partnership under the 2021–2022 BiodivProtect joint call for research proposals, with funding to AGG, MCP and AGG by the ANR Agence Nationale de la Recherche (ANR-22-EBIP-0009-06). MN received funding from the European Union’s Horizon 2020 research and innovation programme under the Marie Skłodowska-Curie grant agreement No 101104374.

## Notes

### Competing Interest Statement

The authors have declared no competing interest.

