## Appendix_Gonzalez_Garcia_et_al_2026 for "Ecosystem service gradients at protected area borders reveal multiple patterns and prevalent management conflicts"

### Ecosystem service gradients vary along protected area borders and across value domains

#### Index

**S1. Study area description**

**S2. Methodological details: Biophysical models**
S2.1. Food production model
S2.2. Outdoor recreation model
S2.3. Carbon storage and sequestration model
S2.4. Heatwave mitigation model
S2.5. Flood regulation model
S2.6. Erosion control model

**S3. Methodological details: Connectivity-based models**
S3.1. General workflow: from species to ecosystem services
S3.2. Species distribution modelling workflow
S3.3. Pollination model
S3.4. Mosquito control model
S3.5. Hunting value model
S3.6. Biological control model
S3.7. Seed dispersal model
S3.8. Emblematic species model

**S4. Methodological details: Structural buffers and sampling protocol**
S4.1. Structural buffers creation
S4.2. Sampling methods details
S4.3. Border typology generation: Algorithmic details

**S5. Methodological details: Statistical analysis of landscape drivers**

**S6. Methodological details: Robustness analysis**
S6.1. Geometric similarity analysis (Correlation)
S6.2. Categorical similarity analysis (Typology agreement)
S6.3. Place-based ecosystem services analysis

**S7. Supporting information for results interpretation**

S7.1. Additional information for ecosystem services interpretation
S7.2. Robustness analysis results
S7.3. Detailed statistical results for landscape drivers
S7.4. The role of place-based ecosystem services in shaping border patterns

**S8. Methodological interpretations and considerations**

**S9. Supplementary references**

#### S1. Study Area Description

Our study was conducted in the Grenoble region, located in the French Alps (Figure S1). This region represents a highly relevant case study for analyzing the interfaces between protected areas and human-modified landscapes. It is characterized by strong altitudinal and environmental gradients, ranging from densely populated and agriculturally intensive valleys (e.g., the Grésivaudan valley where Grenoble city is located) to high-mountain environments.

The landscape is a complex mosaic of land uses, including urban and industrial areas, extensive croplands (such as maize and walnuts), grasslands, and large expanses of forest, covering a total area of approximately 3,800 km². This heterogeneous matrix hosts a diverse network of 16 protected areas, including national parks, regional natural parks, and nature reserves. These protected areas vary greatly in size, protection level, and the socio-economic context of their surroundings, making the region an ideal natural laboratory for studying the diverse patterns of ecosystem services across their borders.

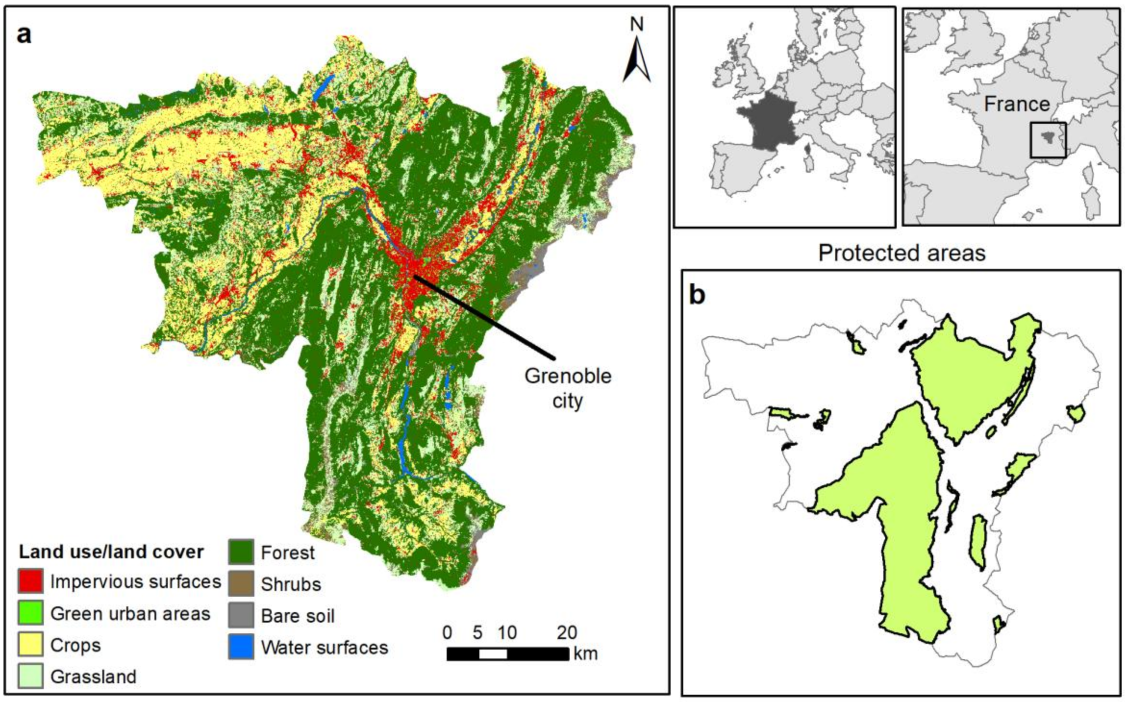


**Figure S 1.** Description of the study area. The panels show the geographical location of the Grenoble region within Europe and France. (a) Land use/land cover map of the study area, illustrating the heterogeneous mosaic of urban centers (e.g., Grenoble city), agricultural valleys, and forested mountain slopes. (b) The location of the 16 protected areas analyzed in this study.

#### S2. Methodological details: Biophysical models

##### S2.1. Food production model

We calculated food production as the energy equivalent value of the average annual crop production based on national estimates (Lasseur et al., 2018). We used national data on yields by crop (Agreste, 2024), provided in 100 kJ/ha, and then calculated the energy value in GJ/ha based on Pérez-Soba et al. (2015) (Table S1). We then calculated the production rate for the 5m pixel resolution of our land use land cover map and created the layer in ArcGIS 10.7. The food production model does not include additional demand or access variables. As food produced in the region is primarily traded on national and international markets, its demand is considered delocalized and not solely tied to the local population, making a supply-focused indicator the most appropriate for this ecosystem service.

**Table S 1.** Yields used for the food production model. "Yield (100 kg/ha)" represents the direct value from public authorities for each crop type. "Tn/ha" is the conversion to tons per hectare. "Energy (MJ/kg)" represents the transformation of yields into energy, and "GJ/5m" is the final conversion based on a 5-meter pixel size.

| CODE | Description | Yield 100kg/ha | Tn/ha | Energy Mj/kg | GJ/5m |
|---|---|---|---|---|---|
| **21000** | Cultivated areas - Arable land - Annual crops | 58.7 | 5.9 | 11.3 | 3.3 |
| **21211** | Common wheat | 54.0 | 5.4 | 11.4 | 3.1 |
| **21212** | Durum wheat | 53.0 | 5.3 | 11.4 | 3.0 |
| **21213** | Barley | 68.0 | 6.8 | 11.5 | 3.9 |
| **21214** | Rye | 44.0 | 4.4 | 12.1 | 2.7 |
| **21215** | Oats | 43.0 | 4.3 | 10.2 | 2.2 |
| **21216** | Maize | 98.0 | 9.8 | 11.0 | 5.4 |
| **21218** | Triticale | 51.0 | 5.1 | 11.4 | 2.9 |
| **21221** | Potatoes | 422.0 | 42.2 | 2.7 | 5.8 |
| **21222** | Sugar beet | 428.0 | 42.8 | 2.4 | 5.1 |
| **21230** | Other non permanent industrial crops | 52.7 | 5.3 | 37.1 | 9.8 |
| **21231** | Sunflower | 25.0 | 2.5 | 15.3 | 1.9 |
| **21232** | Rape and turnip rape | 32.0 | 3.2 | 15.3 | 2.4 |
| **21233** | Soya | 24.0 | 2.4 | 10.2 | 1.2 |
| **21240** | Dry pulses | 28.0 | 2.8 | 14.0 | 2.0 |
| **21250** | Fodder crops (cereals and leguminous) | 138.0 | 13.8 | 2.0 | 1.4 |
| **21290** | Bare arable land | 0.0 | 0.0 | 0.0 | 0.0 |
| **22000** | Permanent crops | 12.0 | 1.2 | 15.6 | 0.9 |
| **22100** | Vinyard | 83.0 | 8.3 | 2.9 | 1.2 |
| **22200** | Orchard | 286.6 | 28.7 | 6.9 | 2.8 |
| **23100** | Managed grassland - Pastures | 83.0 | 8.3 | 3.8 | 1.6 |
| **23200** | Seminatural grassland - Meadows | 58.0 | 5.8 | 3.8 | 1.1 |
| **32100** | Alpine and sub-alpine natural grassland | 146.0 | 14.6 | 3.8 | 2.7 |

#### S2.2. Outdoor recreation model

We created an indicator for the outdoor recreation potential in Grenoble based on the current literature and the most updated land use/land cover data. Previous studies in the region of Grenoble mapped this ecosystem service by applying a mix of indicators such as the degree of nature protection, proximity to aquatic features, scenic value, avoidance of roads and artificial areas, and accessibility criteria (Byczek et al., 2018). To use as much already developed information in the area as possible, we used Byczek et al. (2018) calculations in cases where the data was not linked to land use/land cover, since they developed highly accurate spatial indicators to construct their recreation model. For those indicators that required recalculation based on the new land use/land cover map, we followed a simplified approach based on other studies that have developed similar models (Lavorel et al., 2020). The conceptual design of the model is presented in the Figure S2. To include demand in the model, we introduced an accessibility variable based on crowd-sourced GPS track data, which reflects observed visitor usage patterns and preferences, thereby directly incorporating the spatial dimension of recreational demand.
To calculate each of the variables presented in the Figure S8 we followed different approaches:

- Degree of nature protection: We used the calculation made by Byczek et al. (2018) based in a scoring of the different protected areas in the Grenoble area. The scoring was based on the specific level of conservation (i.e., national park is more restrictive than regional park) and the zones designed in the protected areas plans (i.e., different gradients of protection inside the same protected area), both defined within the French protected area legislation.
- Proximity to aquatic features: We used the land use/land cover map from Marsoner et al. (2023) and extracted the water reservoirs. In the other hand, we used the river network Anon (2021) for the water streams. We created a buffer of 500m of distance from rivers and water bodies to include the proximity effect to aquatic features.
- Proximity to natural features: We used again Marsoner et al. (2023) to calculate this variable. We assigned a value of 1 to those areas linked to natural ecosystems such as forests, alpine grasslands, rivers, etc. We assigned a value of 0.33 to semi-natural areas such as crops and productive grasslands. Finally, we assigned a value of 0 to artificial land uses.
- Scenic value: We used the calculation conducted in Byczek et al. (2018) which consist in a viewshed analyses combined with measures of landscape heterogeneity. In their study they counted the number of pixels in a digital elevation model that can be seen from other pixel in a radius of 500m.
- Road avoidance: We used the variable calculated in Byczek et al. (2018) consisting in the noise avoidance of the road network. They created this indicator based in difference distances for each type of road (i.e., 10m buffer for small roads, 300m for highway and high speed train and 500m from the airport).
- Avoidance of artificialized areas: We used the same layer as for proximity to natural features but in this case we assigned a value of 1 to natural areas such as forests and alpine grasslands or rocky outcrops and 0 value to all artificial and human dominated areas such as crops.
- Accessibility: We used the variable calculated in Byczek et al. (2018) consisting in the use of crowd-sourced websites with information of different tracks for different nature-based activities such as sky, trekking and climbing. This map shows a detailed distribution of the different accessible areas at a very detailed level.

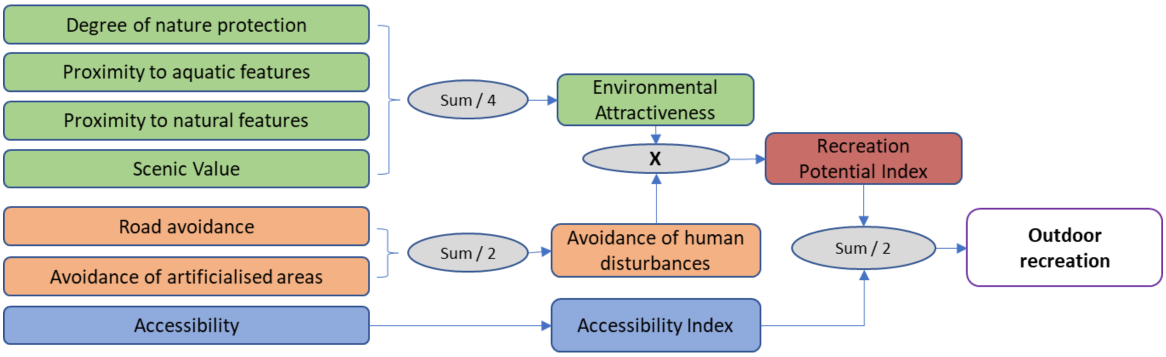


**Figure S 2.** Conceptual design of the outdoor recreation model. From Byczek et al. (2018).

##### S2.3. Carbon storage and sequestration model

We used the InVEST carbon storage model, which consists of a sum of total carbon stored in the four main parts of each ecosystem: carbon in soil, carbon in aboveground biomass, carbon in belowground biomass, and carbon in dead matter. Data was obtained from the literature, focusing on ecosystems with similar characteristics for the Alps. In cases where data was not available, we used data from similar ecosystems in different study sites. Table S2 shows the values used for each land use-land cover category. This model does not include demand or access, as carbon storage is considered an ecosystem service with global beneficiaries, and therefore no specific local demand is required for the analysis.

For urban land use cover, we assumed zero carbon in dead matter (from codes 11000 to 14100) due to the frequent management activities. Carbon in soil for urban parks (code 14100) was considered as the average value of natural ecosystems such as forests, scrubs, and grasslands. Cultivated areas were considered as primarily providing carbon storage in soil, and we did not differentiate between different crops. For permanent crops (code 22000), we used the values for walnut since Grenoble has a significant area of cultivated walnuts. In the case of vineyards (code 22100), we considered the above and below ground carbon the same as for scrubs.

**Table S 2.** Carbon pools in $CO_2$ eq. used for the InVEST model. C_soil is the organic carbon in soil, C_Above is the carbon in aboveground biomass, C_Below is the carbon in belowground biomass, C_litter is the carbon in dead matter.

| CODE | Description | C_soil | C_Above | C_Below | C_litter | References |
|---|---|---|---|---|---|---|
| **11000** | Artificial surfaces and constructions | 0.18 | 0.10 | 0 | 0 | González-García et al. (2020) |
| **11100** | Dense settlement area | 0.18 | 0.10 | 0 | 0 | |
| **11200** | Low density settlement area | 64.04 | 16.86 | 8.57 | 0 | Dorendorf et al. (2015) |
| **11300** | Builtup area | 0.18 | 0.10 | 0 | 0 | González-García et al. (2020) |
| **11400** | Open settlement area | 64.04 | 16.86 | 8.57 | 0.00 | Dorendorf et al. (2015) |
| **12100** | Industrial and commercial zones | 12.34 | 14.44 | 0 | 0 | |
| **12210** | Roads motroways and trunks | 0.18 | 0.10 | 0 | 0 | González-García et al. (2020) |
| **12220** | Road networks | 0.18 | 0.10 | 0 | 0 | |
| **12221** | Roads tertiary and others | 0.18 | 0.10 | 0 | 0 | |
| **12230** | Railways train tracks | 0.18 | 0.10 | 0 | 0 | |

| | | | | | | |
|---|---|---|---|---|---|---|
| **12240** | Unpaved roads and tracks | 0.18 | 0.10 | 0 | 0 | |
| **14100** | Green urban areas | 258.82 | 135.8 | 50.62 | 0 | Dorendorf et al. (2015) |
| **21000** | Cultivated areas - Arable land - Annual crops | 81.68 | 5.07 | 0 | 0 | Dorendorf et al. (2015) |
| **21211** | Common wheat | 81.68 | 5.07 | 0 | 0 | |
| **21212** | Durum wheat | 81.68 | 5.07 | 0 | 0 | |
| **21213** | Barley | 81.68 | 5.07 | 0 | 0 | |
| **21214** | Rye | 81.68 | 5.07 | 0 | 0 | |
| **21215** | Oats | 81.68 | 5.07 | 0 | 0 | |
| **21216** | Maize | 81.68 | 5.07 | 0 | 0 | |
| **21218** | Triticale | 81.68 | 5.07 | 0 | 0 | |
| **21221** | Potatoes | 81.68 | 5.07 | 0 | 0 | |
| **21222** | Sugar beet | 81.68 | 5.07 | 0 | 0 | |
| **21230** | Other non-permanent industrial crops | 81.68 | 5.07 | 0 | 0 | |
| **21231** | Sunflower | 81.68 | 5.07 | 0 | 0 | |
| **21232** | Rape and turnip rape | 81.68 | 5.07 | 0 | 0 | |
| **21233** | Soya | 81.68 | 5.07 | 0 | 0 | |
| **21240** | Dry pulses | 81.68 | 5.07 | 0 | 0 | |
| **21250** | Fodder crops (cereals and leguminous) | 81.68 | 5.07 | 0 | 0 | |
| **21290** | Bare arable land | 81.68 | 0 | 0 | 0 | |
| **22000** | Permanent crops | 151.48 | 52.923 | 20.22 | 0 | Cardinael et al. (2017) |
| **22100** | Vineyard | 153.63 | 6.2 | 8.57 | 0 | Chiti, et al. (2012) |
| **22200** | Orchard | 161.7 | 0 | 0 | 0 | Chiti, et al. (2012) |
| **23100** | Managed grassland - Pastures | 277.97 | 7.13 | 13.2 | 0 | Bahn et al. (2008); Guidi et al. 2014 (managed grassland) |
| **23200** | Seminatural grassland - Meadows | 277.97 | 7.13 | 13.2 | 0 | |
| **31100** | Broadleaf tree cover | 296.75 | 176.41 | 67.42 | 18.11 | INFC 2005, Tonolli & Salvagni, (2007) |
| **31102** | Broadleaf tree cover 30-60% | 296.75 | 52.92 | 20.22 | 5.43 | |
| **31103** | Broadleaf tree cover 60-100% | 296.75 | 105.84 | 40.45 | 10.86 | |
| **31200** | Coniferous tree cover | 258.88 | 267.96 | 76.52 | 30.09 | |

| | | | | | | |
|---|---|---|---|---|---|---|
| **31202** | Coniferous tree cover 30-60% | 258.88 | 80.38 | 22.95 | 9.02 | |
| **31203** | Coniferous tree cover 60-100% | 258.88 | 160.77 | 45.91 | 18.05 | |
| **31300** | Mixed tree cover | 277.81 | 222.18 | 71.97 | 24.1 | |
| **31400** | Tree cover in agricultural context | 151.48 | 52.92 | 20.22 | 0 | Cardinael et al. (2017) |
| **31450** | Tree cover in urban context | 0.18 | 135.8 | 50.62 | 0 | Dorendorf et al. (2015) |
| **31500** | Green linear elements - linear woody features | 84.19 | 6.20 | 8.57 | 4.90 | Sørensen et al., (2018) |
| **31600** | Patchy woody features | 84.19 | 6.20 | 8.57 | 4.90 | |
| **31610** | Additional woody features | 84.19 | 6.20 | 8.57 | 4.90 | |
| **32000** | Scrub and shrubland | 84.19 | 6.20 | 8.57 | 4.90 | |
| **32100** | Alpine and sub-alpine natural grassland | 277.97 | 7.13 | 13.2 | 0 | |
| **32200** | Moors and heathland - other scrubland | 84.19 | 6.20 | 8.57 | 4.90 | |
| **32300** | Sclerophyllous vegetation | 97.8 | 5.69 | 7.5 | 1.53 | Fonseca et al. (2012) |
| **33100** | Beaches, dunes, sands | 18.51 | 0 | 0 | 0 | González-García et al. (2020) |
| **33200** | Bare rocks and rock debris | 0 | 0 | 0 | 0 | |
| **33300** | Sparsely vegetated land | 12.34 | 6.2 | 8.57 | 4.9 | |
| **33500** | Permanent snow covered surfaces | 0 | 0 | 0 | 0 | |
| **41000** | Wetland (permanent wet areas) - inland marshes | 539 | 43 | 0 | 0 | IPCC |
| **51000** | Water bodies | 320.1 | 0.44 | 0.25 | 1.83 | González-García et al. (2020) |
| **51100** | Rivernetwork | 320.1 | 0.44 | 0.25 | 1.83 | |

##### S2.4. Heatwave mitigation model

For the heatwave mitigation ecosystem service, we used the InVEST urban cooling model (Zawadzka et al., 2021). The model estimates the cooling capacity of natural and semi-natural features. This model is primarily designed to assess the cooling effect of green urban areas during a specific heatwave event and its consequent heat island effect. However, the model provides various outputs that can help analyse ecosystem services. Following this approach, we used the cooling capacity as the main reference for the ecosystem service indicator. This model already accounts for demand by integrating the urban heat island effect, which identifies the areas where the cooling effect is most needed. In this case, we used the specific temperature values of Grenoble's heat island effect. Therefore, no additional demand variables were applied.

The variables required for the model include a land use/land cover map and a biophysical table with associated parameters for the different land use/land cover types. This biophysical table includes the crop coefficient value, a true/false value indicating if the land use/land cover is a natural element, the shade ratio, and the albedo. The values used for each land use/land cover type can be found in Table S3 which we obtained from literature analysis.

Additional variables needed for the heatwave mitigation model include the evapotranspiration, obtained from Chelsa (Karger et al., 2021), and the maximum cooling distance, which is a fixed value representing the maximum distance over which a green area larger than 2 hectares has a cooling effect (450m following user guide recommendations). Similarly, the model uses a fixed value for the air blending distance, which we set at 500m following user guide recommendations.

Furthermore, two variables representing the reference temperature and the heat island effect are required. For these, we used a recent study conducted in the Grenoble region (Foissard et al., 2024). The study provides an in-depth analysis of the heat island effect in the region, finding a 4.4 ºC difference between the city and the rural reference temperature by analysing various climate stations. The rural reference temperature was set to 30ºC.

**Table S 3**. Biophysical table associated with the heatwave mitigation model. Albedo represents the albedo of each land use/land cover type. Kc refers to the evapotranspiration coefficients for each land use/land cover type. Green area indicates whether the area is natural or not. Shade represents the shade ratio for each land use/land cover type.

| CODE | Description | Albedo | Kc | Green area | Shade | References |
|---|---|---|---|---|---|---|
| **11000** | Artificial surfaces and constructions | 0.143 | 0.325 | 0 | 0 | Trlica et al., (2017) |
| **11100** | Dense settlement area | 0.143 | 0.325 | 0 | 0 | |
| **11200** | Low density settlement area | 0.147 | 0.25 | 0 | 0 | |
| **11300** | Builtup area | 0.143 | 0.325 | 0 | 0 | |
| **11400** | Open settlement area | 0.147 | 0.25 | 0 | 0 | |
| **12100** | Industrial and commercial zones | 0.132 | 0.35 | 0 | 0 | |
| **12210** | Roads motroways and trunks | 0.139 | 0.3 | 0 | 0 | |
| **12220** | Road networks | 0.139 | 0.3 | 0 | 0 | |
| **12221** | Roads tertiary and others | 0.139 | 0.3 | 0 | 0 | |
| **12230** | Railways train tracks | 0.139 | 0.3 | 0 | 0 | |
| **12240** | Unpaved roads and tracks | 0.132 | 0.31 | 0 | 0 | |
| **14100** | Green urban areas | 0.136 | 0.27 | 1 | 0 | average (31100, 31200, 31300, 32000, 32100) |
| **21000** | Cultivated areas - Arable land - Annual crops | 0.180 | 0.912 | 1 | 0 | Average annual crops |
| **21211** | Common wheat | 0.186 | 0.89 | 1 | 0 | Sieber et al. (2022), Kang et al. (2003) |
| **21212** | Durum wheat | 0.186 | 0.8 | 1 | 0 | Lhomme et al. (2009) |
| **21213** | Barley | 0.176 | 1.07 | 1 | 0 | Sieber et al. (2022), Attarod et al. (2009) |
| **21214** | Rye | 0.181 | 0.87 | 1 | 0 | Sieber et al. (2022), |
| **21215** | Oats | 0.172 | 0.867 | 1 | 0 | Andréasson, (2023), Moteva et al. (2014) |

| | | | | | | |
|---|---|---|---|---|---|---|
| **21216** | Maize | 0.184 | 0.973 | 1 | 0 | Bsaibes et al. (2009), Kang et al. (2003) |
| **21218** | Triticale | 0.183 | 0.978 | 1 | 0 | Moteva et al. (2014) |
| **21221** | Potatoes | 0.214 | 0.65 | 1 | 0 | Paredes et al. (2018) |
| **21222** | Sugar beet | 0.179 | 0.918 | 1 | 0 | Sieber et al. (2022), Moteva et al. (2014) |
| **21230** | Other non-permanent industrial crops | 0.180 | 0.912 | 1 | 0 | 21000 |
| **21231** | Sunflower | 0.225 | 0.79 | 1 | 0 | Srivastava et al. (1998), Mila et al. (2016) |
| **21232** | Rape and turnip rape | 0.188 | 0.945 | 1 | 0 | Sieber et al. (2022), Moteva et al. (2014) |
| **21233** | Soya | 0.239 | 0.86 | 1 | 0 | Weiss et al. (2001), Attarod et al. (2009) |
| **21240** | Dry pulses | 0.214 | 0.8 | 1 | 0 | Nandi et al. (2024) |
| **21250** | Fodder crops (cereals and leguminous) | 0.214 | 0.884 | 1 | 0 | Weiss et al. (2001), Moteva et al. (2014) |
| **21290** | Bare arable land | 0.23 | 0.275 | 1 | 0 | 33100 |
| **22000** | Permanent crops | 0.115 | 1.075 | 1 | 0.7 | Schwaab et al., (2015) |
| **22100** | Vineyard | 0.2 | 0.86 | 1 | 0 | Galleguillos et al. (2011), Cancela et al. (2010) |
| **22200** | Orchard | 0.179 | 0.918 | 1 | 0 | Sieber et al. (2022), Moteva et al. (2014) |
| **23100** | Managed grassland - Pastures | 0.196 | 0.93 | 1 | 0 | Rosset et al. (2001), Ambrosi et al. (2024) |
| **23200** | Seminatural grassland - Meadows | 0.168 | 0.93 | 1 | 0 | |
| **31100** | Broadleaf tree cover | 0.126 | 1.55 | 1 | 1 | Schwaab et al., (2015) |
| **31102** | Broadleaf tree cover 30-60% | 0.126 | 1.55 | 1 | 0.45 | |
| **31103** | Broadleaf tree cover 60-100% | 0.126 | 1.55 | 1 | 0.8 | |
| **31200** | Coniferous tree cover | 0.104 | 1 | 1 | 1 | |
| **31202** | Coniferous tree cover 30-60% | 0.104 | 1 | 1 | 0.45 | |
| **31203** | Coniferous tree cover 60-100% | 0.104 | 1 | 1 | 0.8 | |
| **31300** | Mixed tree cover | 0.115 | 1.4 | 1 | 1 | |
| **31400** | Tree cover in agricultural context | 0.115 | 1.075 | 1 | 0.7 | |
| **31450** | Tree cover in urban context | 0.136 | 0.27 | 1 | 0.7 | 14100 |
| **31500** | Green linear elements - linear woody features | 0.132 | 0.975 | 1 | 0.4 | |
| **31600** | Patchy woody features | 0.132 | 0.975 | 1 | 0 | |

| | | | | | | |
|---|---|---|---|---|---|---|
| **31610** | Additional woody features | 0.132 | 0.975 | 1 | 0 | |
| **32000** | Scrub and shrubland | 0.132 | 0.975 | 1 | 0 | Aartsma et al., (2020) |
| **32100** | Alpine and sub-alpine natural grassland | 0.194 | 1.125 | 1 | 0 | Tian et al., (2014); Blumthaler and Ambach, (1988) |
| **32200** | Moors and heathland - other scrubland | 0.132 | 0.975 | 1 | 0 | Aartsma et al., (2020) |
| **32300** | Sclerophyllous vegetation | 0.132 | 0.65 | 1 | 0 | |
| **33100** | Beaches, dunes, sands | 0.23 | 0.275 | 1 | 0 | Blumthaler and Ambach, (1988) |
| **33200** | Bare rocks and rock debris | 0.3 | 0.125 | 1 | 0 | Al Fahdawi, (2021) |
| **33300** | Sparsely vegetated land | 0.13 | 0.55 | 1 | 0 | Sieber et al., (2022) |
| **33500** | Permanent snow covered surfaces | 0.507 | 0.95 | 1 | 0 | Kalitin, (1930) |
| **41000** | Wetland (permanent wet areas) - inland marshes | 0.159 | 0.625 | 1 | 0 | Trlica et al., (2017) |
| **51000** | Water bodies | 0.091 | 0.95 | 1 | 0 | |
| **51100** | Rivernetwork | 0.091 | 0.95 | 1 | 0 | Blumthaler and Ambach, (1988) |
| **51200** | Riverbed > 10m width | 0.091 | 0.95 | 1 | 0 | |

##### S2.5. Flood regulation model

For the flood regulation ecosystem service, we used the InVEST urban flood regulation model, which calculates the avoided runoff for each pixel based on the land use/land cover type and the curve numbers for these covers in each soil hydrologic group. The land use/land cover layer needs to be associated with a biophysical table that includes the curve number value (Table S4). For the soil hydrologic group, we used the Ross et al. (2018) database, which includes the four main types of soil hydrologic groups for Europe at a 250m pixel resolution. The model also uses a fixed value representing a reference rain event to calculate the total water retained by ecosystems; we used a fixed value of 100 mm for this parameter. The output of the model includes various variables, such as the runoff retention index and the amount of water retained in cubic meters ($m^3$). For this study, we used the runoff retention index to capture the potential capacity of ecosystems to retain water. We did not include additional changes related to demand, as we considered that the selected reference precipitation of 100mm, representing a high-intensity storm event, already reflects the societal demand for flood regulation under risk scenarios.

**Table S 4.** Biophysical table associated with the flood regulation model. CN values represent the curve numbers for each soil hydrologic group (A, B, C, or D). The "Ref" field indicates the equivalencies used from the main sources related to land use/land cover categories.

| CODE | Description | CN_a | CN_b | CN_c | CN_d | Reference | Ref |
|---|---|---|---|---|---|---|---|
| **11000** | Artificial surfaces and constructions | 98 | 98 | 98 | 98 | | Impervious areas |
| **11100** | Dense settlement area | 98 | 98 | 98 | 98 | | |

| | | | | | | | |
|---|---|---|---|---|---|---|---|
| **11200** | Low density settlement area | 61 | 75 | 83 | 87 | | 1⁄4 acre (38% imp.) |
| **11300** | Builtup area | 98 | 98 | 98 | 98 | | Impervious areas |
| **11400** | Open settlement area | 61 | 75 | 83 | 87 | Cronshey (1986) | 1⁄4 acre (38% imp.) |
| **12100** | Industrial and commercial zones | 98 | 98 | 98 | 98 | | Impervious areas |
| **12210** | Roads motroways and trunks | 98 | 98 | 98 | 98 | | |
| **12220** | Road networks | 98 | 98 | 98 | 98 | | |
| **12221** | Roads tertiary and others | 98 | 98 | 98 | 98 | | |
| **12230** | Railways train tracks | 98 | 98 | 98 | 98 | | |
| **12240** | Unpaved roads and tracks | 77 | 86 | 91 | 94 | | Bare soil |
| **14100** | Green urban areas | 39 | 61 | 74 | 80 | | Open space (Good condition) |
| **21000** | Cultivated areas - Arable land - Annual crops | 60 | 72 | 80 | 84 | | Small grain SR+CR good |
| **21211** | Common wheat | 60 | 72 | 80 | 84 | | |
| **21212** | Durum wheat | 60 | 72 | 80 | 84 | | |
| **21213** | Barley | 60 | 72 | 80 | 84 | | |
| **21214** | Rye | 60 | 72 | 80 | 84 | | |
| **21215** | Oats | 60 | 72 | 80 | 84 | | |
| **21216** | Maize | 65 | 75 | 82 | 86 | | Row crops contoured |
| **21218** | Triticale | 60 | 72 | 80 | 84 | | Small grain SR+CR good |
| **21221** | Potatoes | 32 | 58 | 72 | 79 | | Orchard |
| **21222** | Sugar beet | 32 | 58 | 72 | 79 | | |
| **21230** | Other non permanent industrial crops | 60 | 72 | 80 | 84 | | Small grain SR+CR good |
| **21231** | Sunflower | 64 | 75 | 82 | 85 | | Row crops SR+CR |
| **21232** | Rape and turnip rape | 64 | 75 | 82 | 85 | | |

| | | | | | | | |
|---|---|---|---|---|---|---|---|
| **21233** | Soya | 58 | 72 | 81 | 85 | | Close-seeded or broadcast legumes or rotation meadow |
| **21240** | Dry pulses | 58 | 72 | 81 | 85 | Cronshey (1986) | |
| **21250** | Fodder crops (cereals and leguminous) | 58 | 72 | 81 | 85 | | |
| **21290** | Bare arable land | 77 | 86 | 91 | 94 | | Bare arable land |
| **22000** | Permanent crops | 43 | 65 | 76 | 82 | | Agro-forestry areas |
| **22100** | Vinyard | 67 | 78 | 85 | 89 | | Straight row |
| **22200** | Orchard | 32 | 58 | 72 | 79 | | Orchard |
| **23100** | Managed grassland - Pastures | 8 | 79 | 86 | 89 | | Pasture |
| **23200** | Seminatural grassland - Meadows | 30 | 58 | 71 | 78 | | Meadow |
| **31100** | Broadleaf tree cover | 30 | 55 | 70 | 77 | | Good condition |
| **31102** | Broadleaf tree cover 30-60% | 36 | 60 | 73 | 79 | | Fair condition |
| **31103** | Broadleaf tree cover 60-100% | 30 | 55 | 70 | 77 | | Good condition |
| **31200** | Coniferous tree cover | 33 | 58 | 72 | 78 | | Good condition |
| **31202** | Coniferous tree cover 30-60% | 40 | 63 | 75 | 80 | Jaafar et al. (2019); Tedela et al. (2012) | Fair condition |
| **31203** | Coniferous tree cover 60-100% | 33 | 58 | 72 | 78 | | Good condition |
| **31300** | Mixed tree cover | 31.5 | 56.5 | 71 | 77.5 | | Average broadleaf and conifer |
| **31400** | Tree cover in agricultural context | 43 | 65 | 76 | 82 | Cronshey (1986) | Woods—grass combination (orchard or tree farm).D (Fair) |
| **31450** | Tree cover in urban context | 39 | 61 | 74 | 80 | | Open space (Good condition) |
| **31500** | Green linear elements - linear woody features | 43 | 65 | 76 | 82 | | Woods—grass combination (orchard or tree farm).D |
| **31600** | Patchy woody features | 43 | 65 | 76 | 82 | | |

| | | | | | | | |
|---|---|---|---|---|---|---|---|
| **31610** | Additional woody features | 43 | 65 | 76 | 82 | | |
| **32000** | Scrub and shrubland | 43 | 65 | 76 | 82 | | |
| **32100** | Alpine and sub-alpine natural grassland | 39 | 61 | 74 | 80 | | Brush—brush-weed-grass mixture with<br><br>brush the major element.B (Fair) |
| **32200** | Moors and heathland - other scrubland | 35 | 56 | 70 | 77 | | Brush—brush-weed-grass mixture with brush the major element.B |
| **32300** | Sclerophyllous vegetation | 0 | 62 | 74 | 85 | | Herbaceosu for arid rangelands |
| **33100** | Beaches, dunes, sands | 63 | 77 | 85 | 88 | | Natural desert landscaping (pervious areas only) |
| **33200** | Bare rocks and rock debris | 63 | 77 | 85 | 88 | | |
| **33300** | Sparsely vegetated land | 68 | 79 | 86 | 89 | | Poor condition (grass cover < 50%) |
| **33500** | Permanent snow covered surfaces | 99 | 99 | 99 | 99 | | |
| **41000** | Wetland (permanent wet areas) - inland marshes | 49 | 69 | 79 | 84 | Chen et al. (2014) | Wet tussock grassland with herbs, sedges or rushes, herblands or ferns |
| **51000** | Water bodies | 99 | 99 | 99 | 99 | | |
| **51100** | Rivernetwork | 99 | 99 | 99 | 99 | | |
| **51200** | Riverbed > 10m width | 99 | 99 | 99 | 99 | | |

##### S2.6. Erosion control model

The Sediment Delivery Ratio model, implemented within the InVEST software suite, is designed to quantify sediment export from watersheds and assess the role of terrestrial vegetation in sediment retention. This model employs an empirical approach to estimate the fraction of soil erosion that is effectively delivered to the watershed outlet. The Sediment Delivery Ratio model operates on a spatially explicit basis, integrating key biophysical factors influencing soil erosion and transport. These factors include rainfall erosivity (R) and soil erodibility (K), topography (LS factor), and land cover/management (C factor), closely mirroring the Revised Universal Soil Loss Equation (RUSLE) framework. Rainfall erosivity (R) and soil erodibility (K) data were obtained from ESDAC (Panagos, et al., 2014, 2017) at resolutions of 1km and 500m, respectively.

Sediment Delivery Ratio calculation is performed at the pixel level, considering hydrological flow paths derived from a Digital Elevation Model. The digital elevation model used has a 5m resolution and was downloaded from the BD Alti for Isère (IGN, 2023), with tiles merged using the merge function in QGIS. A threshold flow accumulation of 400 pixels was used in InVEST to delineate flow paths. The model computes sediment retention capacity along these flow paths, primarily attributed to land cover characteristics. Land use data, at a 5m resolution, was sourced from the EUSALP map (Marsoner et al., 2023). Areas with higher vegetation cover are assumed to exhibit greater sediment interception and retention potential. By integrating these factors, and using parameters such as a Borselli K value of 2 and a Borselli IC0 of 0.8 within the model, the Sediment Delivery Ratio model estimates the total eroded sediment at each raster cell and the proportion of this sediment that is retained or transported downstream. This process culminates in a spatially distributed estimation of Sediment Delivery Ratio, constrained by a maximum sediment delivery ratio value of 0.5, and the total sediment load exported from the watershed. A maximum L value of 122 was used for the LS factor calculation within the model.

Demand for erosion control is implicitly included in this model. The model calculates sediment export based on rainfall erosivity (R factor), which represents the risk of erosion. The ecosystem service is therefore inherently higher in areas where this risk, and thus the need for control, is greatest. We did not add further demand layers, as the ecosystem service is relevant across all non-hardened surfaces where soil loss can occur.

#### S3. Methodological details: Connectivity-based models

##### S3.1. General workflow: from species to ecosystem services

To model ecosystem services provided by mobile organisms, we selected a total of 224 vertebrate species present in the French Alps. The primary criterion for inclusion was the availability of sufficient occurrence data to build robust species distribution models. These species were then assigned to one or more of the six connectivity-dependent ecosystem services based on their ecological traits and functions. For the aggregation of species-level maps, we applied an equal weighting to all species contributing to a given ecosystem service. While individual species undoubtedly vary in their functional efficiency, quantifying these differences for over 200 species was beyond the scope of this study. This averaging approach, therefore, represents a robust and common method for creating a composite index of the overall potential for a given ecosystem service, assuming a principle of functional redundancy where the presence of a diverse assemblage is more important than the contribution of any single species.

The rationale for each ecosystem service grouping is as follows:

- Hunting Value: This group includes 24 species that are officially listed as hunting species in the region, based on data provided by regional hunters' associations.
- Biological Control: This group comprises 78 species known to prey on agricultural pests, including various insectivorous birds, bats, and small mammals.
- Mosquito Control: This group consists of 81 species, primarily bats and birds, that are known predators of mosquitoes and other nuisance insects.
- Seed Dispersal: This group includes 81 species that play a key role in the dispersal of native plant species, contributing to forest regeneration and ecosystem resilience.
- Emblematic Species: This group is composed of 96 species selected for their high cultural and touristic value. The selection criteria included species of community interest under European legislation (e.g., Annexes of the Habitats and Birds directives), species frequently mentioned on regional tourism websites, and species with high observation rates relative to their range size, indicating significant public interest.
- Pollination: Due to the complexity of modeling hundreds of pollinator species, we used a pollinator archetype approach. This involves creating a single functional profile representing a typical pollinator in the region, defined by characteristics such as median foraging distance and habitat preferences, based on the extensive literature on wild bees and other insect pollinators.

A complete list detailing the assignment of each of the 224 vertebrate species to the different ecosystem services can be found in Table S5.

**Table S 5.** Species used for each ecosystem service based on connectivity algorithms. "Total ES" indicates the number of times a species appears in different ecosystem services.

| Species | Hunting value | Biological control | Seed dispersal | Mosquito control | Emblematic species | Total ES |
|---|---|---|---|---|---|---|
| ***Acanthis hornemanni*** | | | x | | | 1 |
| ***Accipiter gentilis*** | | x | | | x | 2 |
| ***Accipiter nisus*** | | | | | x | 1 |
| ***Aegolius funereus*** | | x | | x | x | 3 |
| ***Aegypius monachus*** | | | | | x | 1 |
| ***Alauda arvensis*** | | x | | | | 1 |
| ***Alcedo atthis*** | | | | | x | 1 |
| ***Alectoris graeca*** | | | x | | x | 2 |
| ***Alectoris rufa*** | x | | x | | | 2 |
| ***Alytes obstetricans*** | | x | | | x | 2 |
| ***Anas platyrhynchos*** | x | | x | | | 2 |
| ***Anthus campestris*** | | | | | x | 1 |
| ***Anthus pratensis*** | | x | | | | 1 |
| ***Anthus spinoletta*** | | | x | | | 1 |
| ***Apodemus alpicola*** | | | x | | | 1 |
| ***Apodemus flavicollis*** | | | x | | | 1 |
| ***Apodemus sylvaticus*** | | | x | | | 1 |
| ***Apus apus*** | | x | | x | | 2 |
| ***Apus melba*** | | x | | x | | 2 |
| ***Aquila chrysaetos*** | | x | | | x | 2 |
| ***Aquila fasciata*** | | | | | x | 1 |
| ***Ardea purpurea*** | | | | | x | 1 |
| ***Asio otus*** | | x | | | | 1 |
| ***Athene noctua*** | | x | | x | | 2 |
| ***Aythya ferina*** | x | | | | | 1 |
| ***Barbastella barbastellus*** | | | | x | x | 2 |
| ***Bombina variegata*** | | | | x | x | 2 |
| ***Botaurus stellaris*** | | | | | x | 1 |
| ***Bubo bubo*** | | x | | | x | 2 |

| | | | | | | |
|---|---|---|---|---|---|---|
| ***Burhinus oedicnemus*** | | x | | | x | 2 |
| ***Buteo buteo*** | | x | | x | | 2 |
| ***Canis lupus*** | | x | | | x | 2 |
| ***Capra ibex*** | | | | | x | 1 |
| ***Capreolus capreolus*** | x | | | | x | 2 |
| ***Caprimulgus europaeus*** | | x | | x | x | 3 |
| ***Carduelis carduelis*** | | | x | | | 1 |
| ***Castor fiber*** | | | | | x | 1 |
| ***Cecropis daurica*** | | x | | x | | 2 |
| ***Certhia brachydactyla*** | | | | | x | 1 |
| ***Cervus elaphus*** | x | | x | | x | 3 |
| ***Chloris chloris*** | | | x | | | 1 |
| ***Chroicocephalus ridibundus*** | | | | x | | 1 |
| ***Circaetus gallicus*** | | | | | x | 1 |
| ***Circus aeruginosus*** | | x | | | x | 2 |
| ***Circus cyaneus*** | | x | | | x | 2 |
| ***Circus pygargus*** | | x | | | x | 2 |
| ***Columba livia*** | x | | x | | | 2 |
| ***Columba oenas*** | x | | x | | | 2 |
| ***Columba palumbus*** | x | | x | | x | 3 |
| ***Coronella austriaca*** | | | | | x | 1 |
| ***Corvus corax*** | | x | x | x | | 3 |
| ***Corvus corone*** | | x | x | | | 2 |
| ***Corvus frugilegus*** | | x | x | | | 2 |
| ***Corvus monedula*** | | x | x | | | 2 |
| ***Coturnix coturnix*** | x | | x | | | 2 |
| ***Crex crex*** | | | | | x | 1 |
| ***Crocidura russula*** | | x | | | | 1 |
| ***Crocidura suaveolens*** | | x | | | | 1 |
| ***Cuculus canorus*** | | x | | x | | 2 |
| ***Cyanistes caeruleus*** | | | x | | | 1 |
| ***Dama dama*** | x | | | | x | 2 |

| | | | | | |
|---|---|---|---|---|---|
| ***Delichon urbicum*** | x | | x | | 2 |
| ***Dendrocopos major*** | | x | | x | 2 |
| ***Dendrocoptes medius*** | | x | | | 1 |
| ***Dryocopus martius*** | | | | x | 1 |
| ***Egretta garzetta*** | | | | x | 1 |
| ***Eliomys quercinus*** | | x | | | 1 |
| ***Emberiza calandra*** | x | | | | 1 |
| ***Emberiza cia*** | | | x | | 1 |
| ***Emberiza cirlus*** | | x | | | 1 |
| ***Emberiza citrinella*** | x | | | | 1 |
| ***Emberiza hortulana*** | x | | x | x | 3 |
| ***Emberiza schoeniclus*** | | x | x | | 2 |
| ***Emys orbicularis*** | | | | x | 1 |
| ***Epidalea calamita*** | x | | | | 1 |
| ***Eptesicus nilssonii*** | | | x | x | 2 |
| ***Eptesicus serotinus*** | | | x | x | 2 |
| ***Erinaceus europaeus*** | x | | | | 1 |
| ***Erithacus rubecula*** | | x | | | 1 |
| ***Falco peregrinus*** | | | | x | 1 |
| ***Falco subbuteo*** | x | | x | | 2 |
| ***Falco tinnunculus*** | x | | | | 1 |
| ***Felis silvestris*** | x | | | x | 2 |
| ***Ficedula hypoleuca*** | | | x | | 1 |
| ***Gallinula chloropus*** | | x | | | 1 |
| ***Garrulus glandarius*** | | x | x | | 2 |
| ***Glaucidium passerinum*** | x | | | x | 2 |
| ***Glis glis*** | x | x | | | 2 |
| ***Gyps fulvus*** | | | | x | 1 |
| ***Hierophis viridiflavus*** | x | | | | 1 |
| ***Himantopus himantopus*** | | | | x | 1 |
| ***Hippolais polyglotta*** | | | x | | 1 |
| ***Hirundo rustica*** | | | x | | 1 |

| | | | | | | |
|---|---|---|---|---|---|---|
| ***Hyla arborea*** | | | | | x | 1 |
| ***Hypsugo savii*** | | | | x | x | 2 |
| ***Ixobrychus minutus*** | | | | | x | 1 |
| ***Jynx torquilla*** | | | x | | | 1 |
| ***Lacerta agilis*** | | x | | | x | 2 |
| ***Lagopus muta*** | | | x | | x | 2 |
| ***Lanius collurio*** | | x | | x | x | 3 |
| ***Lanius excubitor*** | | x | | x | | 2 |
| ***Lanius senator*** | | x | | x | | 2 |
| ***Larus michahellis*** | | x | | x | | 2 |
| ***Lepus europaeus*** | x | | x | | | 2 |
| ***Lepus timidus*** | x | | | | | 1 |
| ***Linaria cannabina*** | | | x | | | 1 |
| ***Lissotriton vulgaris*** | | | | x | x | 2 |
| ***Lullula arborea*** | | x | | | x | 2 |
| ***Luscinia megarhynchos*** | | | x | | | 1 |
| ***Luscinia svecica*** | | | | x | x | 2 |
| ***Lutra lutra*** | | x | | | x | 2 |
| ***Lynx lynx*** | | x | | | x | 2 |
| ***Marmota marmota*** | | | | | x | 1 |
| ***Martes foina*** | | x | x | | | 2 |
| ***Martes martes*** | | x | | | | 1 |
| ***Meles meles*** | | x | x | | | 2 |
| ***Merops apiaster*** | | x | | x | x | 3 |
| ***Micromys minutus*** | | | x | | | 1 |
| ***Microtus agrestis*** | | | x | | | 1 |
| ***Microtus arvalis*** | | | x | | | 1 |
| ***Microtus duodecimcostatus*** | | | x | | | 1 |
| ***Milvus migrans*** | | x | | x | x | 3 |
| ***Milvus milvus*** | | x | | x | x | 3 |
| ***Miniopterus schreibersii*** | | | | x | x | 2 |
| ***Monticola saxatilis*** | | | | x | | 1 |

| | | | | | | |
|---|---|---|---|---|---|---|
| ***Montifringilla nivalis*** | | | | x | | 1 |
| ***Motacilla alba*** | | | | x | | 1 |
| ***Motacilla cinerea*** | | | | x | | 1 |
| ***Motacilla flava*** | | | | x | | 1 |
| ***Muscardinus avellanarius*** | | | x | | x | 2 |
| ***Muscicapa striata*** | | | | x | | 1 |
| ***Mustela erminea*** | | x | | | | 1 |
| ***Mustela nivalis*** | | x | | | | 1 |
| ***Mustela putorius*** | | x | | | | 1 |
| ***Myotis alcathoe*** | | | | x | x | 2 |
| ***Myotis bechsteinii*** | | | | x | x | 2 |
| ***Myotis blythii*** | | | | x | x | 2 |
| ***Myotis brandtii*** | | | | x | x | 2 |
| ***Myotis capaccinii*** | | | | x | x | 2 |
| ***Myotis daubentonii*** | | | | x | x | 2 |
| ***Myotis emarginatus*** | | | | x | x | 2 |
| ***Myotis myotis*** | | | | | x | 1 |
| ***Myotis mystacinus*** | | | | x | x | 2 |
| ***Myotis nattereri*** | | | | | x | 1 |
| ***Nucifraga caryocatactes*** | | | x | x | | 2 |
| ***Numenius arquata*** | | | x | | | 1 |
| ***Nyctalus lasiopterus*** | | | | x | x | 2 |
| ***Nyctalus leisleri*** | | | | x | x | 2 |
| ***Nyctalus noctula*** | | | | x | x | 2 |
| ***Nycticorax nycticorax*** | | x | | | x | 2 |
| ***Oenanthe oenanthe*** | | x | | x | | 2 |
| ***Oriolus oriolus*** | | | x | | | 1 |
| ***Oryctolagus cuniculus*** | x | | x | | | 2 |
| ***Otus scops*** | | x | | x | | 2 |
| ***Parus major*** | | x | x | | | 2 |
| ***Passer domesticus*** | | | x | | | 1 |
| ***Passer montanus*** | | | x | x | | 2 |

| | | | | | | |
|---|---|---|---|---|---|---|
| ***Perdix perdix*** | x | | x | | x | 3 |
| ***Periparus ater*** | | | x | x | | 2 |
| ***Pernis apivorus*** | | x | | x | x | 3 |
| ***Petronia petronia*** | | | x | | | 1 |
| ***Phasianus colchicus*** | x | | x | | | 2 |
| ***Phoenicurus ochruros*** | | | x | x | | 2 |
| ***Phoenicurus phoenicurus*** | | | x | x | | 2 |
| ***Phylloscopus bonelli*** | | | | x | | 1 |
| ***Phylloscopus sibilatrix*** | | | x | | | 1 |
| ***Phylloscopus trochilus*** | | | x | | | 1 |
| ***Pica pica*** | | x | x | x | | 3 |
| ***Pipistrellus kuhlii*** | | | | x | x | 2 |
| ***Pipistrellus nathusii*** | | | | x | x | 2 |
| ***Pipistrellus pipistrellus*** | | | | x | x | 2 |
| ***Plecotus auritus*** | | | | x | x | 2 |
| ***Plecotus austriacus*** | | | | x | x | 2 |
| ***Podarcis muralis*** | | | | | x | 1 |
| ***Poecile montanus*** | | | x | x | | 2 |
| ***Porzana porzana*** | | | | | x | 1 |
| ***Ptyonoprogne rupestris*** | | | | x | | 1 |
| ***Pyrrhocorax graculus*** | | | x | | | 1 |
| ***Pyrrhocorax pyrrhocorax*** | | | x | | x | 2 |
| ***Pyrrhula pyrrhula*** | | | x | | | 1 |
| ***Rallus aquaticus*** | | | x | | | 1 |
| ***Rana dalmatina*** | | | | | x | 1 |
| ***Rhinolophus euryale*** | | | | x | x | 2 |
| ***Rhinolophus ferrumequinum*** | | | | x | x | 2 |
| ***Rhinolophus hipposideros*** | | | | x | x | 2 |
| ***Riparia riparia*** | | | | x | | 1 |
| ***Rupicapra rupicapra*** | x | | | | x | 2 |
| ***Saxicola rubetra*** | | x | x | | | 2 |
| ***Sciurus vulgaris*** | | | x | | | 1 |

| | | | | | | |
|---|---|---|---|---|---|---|
| ***Scolopax rusticola*** | x | | x | | | 2 |
| ***Sitta europaea*** | | | x | | | 1 |
| ***Sorex araneus*** | | x | | | | 1 |
| ***Sorex coronatus*** | | x | | | | 1 |
| ***Spatula clypeata*** | x | | | | | 1 |
| ***Spatula querquedula*** | x | | | | | 1 |
| ***Spinus spinus*** | | | x | | | 1 |
| ***Streptopelia decaocto*** | x | | x | | | 2 |
| ***Streptopelia turtur*** | x | | x | | | 2 |
| ***Strix aluco*** | | x | | x | | 2 |
| ***Sturnus vulgaris*** | | x | x | | | 2 |
| ***Sus scrofa*** | x | | x | | x | 3 |
| ***Sylvia atricapilla*** | | | | x | | 1 |
| ***Sylvia borin*** | | | x | x | | 2 |
| ***Sylvia communis*** | | | x | | | 1 |
| ***Tadarida teniotis*** | | | | x | x | 2 |
| ***Talpa europaea*** | | x | | | | 1 |
| ***Tetrao tetrix*** | x | | x | | | 2 |
| ***Tetrastes bonasia*** | x | | x | | | 2 |
| ***Tetrax tetrax*** | | | x | | x | 2 |
| ***Timon lepidus*** | | x | | | | 1 |
| ***Triturus cristatus*** | | | | x | x | 2 |
| ***Troglodytes troglodytes*** | | | x | x | x | 3 |
| ***Turdus merula*** | | x | | | | 1 |
| ***Turdus philomelos*** | | | x | | | 1 |
| ***Turdus pilaris*** | | x | x | x | | 3 |
| ***Turdus torquatus*** | | | x | | | 1 |
| ***Turdus viscivorus*** | | x | x | x | | 3 |
| ***Tyto alba*** | | x | | | | 1 |
| ***Upupa epops*** | | x | | | | 1 |
| ***Vanellus vanellus*** | | x | | | | 1 |
| ***Vespertilio murinus*** | | | | | x | 1 |

| | | | | | |
|---|---|---|---|---|---|
| ***Vespertilio murinus.*** | | | x | | 1 |
| ***Vipera aspis*** | x | | | | 1 |
| ***Vulpes vulpes*** | x | x | | | 2 |
| ***Zamenis longissimus*** | x | | | | 1 |
| ***Zootoca vivipara*** | | | | x | 1 |
| ***Zootoca vivipara.*** | x | | | | 1 |

##### S3.2. Species distribution modelling workflow

Habitat suitability modelling (1km European scale):
Species-specific habitat suitability models were developed at 1 km² resolution for terrestrial vertebrates across Europe using an ensemble species distribution modeling approach.

The calibration domain extended beyond the EEA39 reporting region (EU27, the United Kingdom, the four EFTA countries—Iceland, Liechtenstein, Norway, Switzerland—and Turkey) to include adjacent areas in Russia, North Africa, and the Middle East to reduce niche truncation (Thuiller et al., 2004; Barbet-Massin et al., 2010).

- Environmental predictors: The ensemble species distribution models were built using a comprehensive set of environmental predictor variables relevant to species distributions at the European scale and assembled at 1 km² resolution from publicly available datasets. Annual bioclimatic variables averaged over the period 1990-2020 including mean temperature, total precipitation, growing degree days, temperature and precipitation seasonality as well as snow cover days were obtained from CHELSA (Karger et al., 2017; Karger et al., 2020). Topographic variables (slope, landforms, coastal proximity) were derived from EarthEnv (Amatulli et al., 2018). Soil physical and chemical properties (pH, soil organic carbon, texture) were taken from SoilGrids (Poggio et al., 2021). Hydrographic layers describing the binary presence and salinity of water bodies and wetlands were compiled from Corine land cover (Copernicus Land Monitoring Service, 2019) and the Global Lake and Wetland database (Lehner & Döll, 2004), while river density was computed using the EU-Hydro (EEA, 2019) and HydroSHEDS (Lehner et al., 2008) watercourses polygons. Land systems and land-use intensity were obtained from Sandström et al. (2023). To minimise collinearity, only variables with pairwise correlation < 0.70 were retained.

- Presence/absence data: Given the presence-only nature of the source data, we developed a robust process for data preparation. Presence data for each of the 224 vertebrate species were downloaded and curated from the Global Biodiversity Information Facility (GBIF)[1] over the period 1990-2022. To train the models, two types of synthetic absence data were generated: pseudo-absences sampled within species' known ranges using a "target group strategy," and background points sampled outside known ranges to define the environmental space (Phillips et al., 2009).

- Ensemble modeling workflow: For each species, we used five independent datasets (with different random samples of pseudo-absence/background points) and partitioned each into five folds using spatial block cross-validation to ensure spatial independence between training and validation data (Roberts et al., 2017). For each fold, we fitted three

---

[1] Aves : https://doi.org/10.15468/dl.6ptbvk ; https://doi.org/10.15468/dl.6am8ch ; https://doi.org/10.15468/dl.6q3zpq ; https://doi.org/10.15468/dl.y8nfqw ; https://doi.org/10.15468/dl.gdta5n ; https://doi.org/10.15468/dl.tn2b3w ;
Amphibia/Mammalia/Reptilia: https://doi.org/10.15468/dl.rxfzz3

machine-learning algorithms: Random Forest, XGBoost, and Neural Networks. This resulted in an ensemble of 75 fitted models per species. Only individual models achieving a cross-validated True Skill Statistic (TSS) score above a threshold of 0.4 were retained for the final ensemble average. The final ensemble habitat suitability map for each species represents a consensus prediction from multiple algorithms and data partitions, ensuring high robustness (Si-moussi & Thuiller, 2024).

Downscaling of suitability maps to 5m resolution:

To bridge the scale gap between the coarse European habitat suitability predictions (1km) and the fine-scale landscape of our study area (5m), we implemented a multi-step downscaling procedure. This process refines the coarse predictions by integrating them with high-resolution, local land-cover information.

- Species habitat preferences: We first established a correspondence between each species' habitat requirements and the land-cover classes defined in the regional high resolution land cover (EUSALP, Marsoner et al., 2023). Species–habitat associations were based on documented habitat preferences from O'Connor et al. (2024), originally defined for GlobCover classes. To align these with the EUSALP classification, we constructed a crosswalk table linking each EUSALP class to its corresponding GlobCover class. This allowed us to infer species-specific habitat suitability for EUSALP classes, designating each class as suitable or unsuitable for every species (Table S6).
- Binary habitat mask creation (5m): Based on the species–habitat preferences described above, we generated a binary habitat mask at 5 m resolution for each species. Each pixel was assigned a value of 1 if its EUSALP land-cover class was classified as suitable habitat for the species, and 0 otherwise. These masks delineate the spatial distribution of potentially suitable local habitats at fine spatial resolution.
- To integrate fine- and coarse-scale information, we first resampled the European 1 km habitat-suitability map for each species to 5 m resolution using bilinear interpolation, thereby downscaling the broad environmental suitability gradient. The resampled 5 m suitability map was then multiplied by the corresponding 5 m binary habitat mask. This operation filtered broad-scale suitability predictions, retaining values only in pixels corresponding to locally suitable habitat. The resulting high-resolution maps thus combine continental-scale climatic suitability with Alps-specific land-cover constraints, providing spatially explicit inputs for subsequent connectivity analyses.

**Table S 6.** Crosswalk references between EUSALP land use cover used for the study and GlobCover.

| Code | EUSALP | GlobCover |
|---|---|---|
| **11000** | Artificial surfaces and constructions | 190 |
| **11100** | Dense settlement area | 190 |
| **11200** | Low density settlement area | 190 |
| **11300** | Builtup area | 190 |
| **11400** | Open settlement area | 190 |
| **12100** | Industrial and commercial zones | 190 |
| **12210** | Roads motorways and trunks | 190 |
| **12220** | Roads primary and secondary | 190 |
| **12221** | Roads tertiary and others | 190 |

| | | |
|---|---|---|
| **12230** | Railways train tracks | 190 |
| **12240** | Unpaved Roads and Tracks | 190 |
| **14100** | Green urban areas | 190 |
| **21000** | Cultivated areas - Arable Land - Annual Crops | 10,11,13,14,15 |
| **21211** | Common wheat | 10,15 |
| **21212** | Durum wheat | 10,15 |
| **21213** | Barley | 10,15 |
| **21214** | Rye | 10,15 |
| **21215** | Oats | 10,15 |
| **21216** | Maize | 10,13 |
| **21217** | Rice | 10,11 |
| **21218** | Triticale | 10,15 |
| **21219** | Other cereals | 10,15 |
| **21221** | Potatoes | 10,14 |
| **21222** | Sugar beet | 10,14 |
| **21223** | Other root crops | 10,14 |
| **21230** | Other non permanent industrial crops | 10,14,15 |
| **21231** | Sunflower | 10,14 |
| **21232** | Rape and turnip rape | 10,14 |
| **21233** | Soya | 10,15 |
| **21240** | Dry pulses | 10,14,15 |
| **21250** | Fodder crops (cereals and leguminous) | 10,14,15 |
| **21290** | Bare arable land | 10,14,15 |
| **22000** | Permanent Crops | 10,16 |
| **22100** | Vinyard | 10,16 |
| **22200** | Orchard | 10,16 |
| **23100** | Managed grassland - Pastures | 20, 21, 30 |
| **23200** | Seminatural grassland - Meadows | 140,141,144,151,110,120 |
| **31100** | Broadleaf tree cover | 40, 41, 50, 60, 100, 101, 32 |
| **31102** | Broadleaf tree cover 30-60% | 40, 41, 50, 60, 100, 101, 32 |
| **31103** | Broadleaf tree cover 60-100% | 40, 41, 50, 60, 100, 101, 32 |
| **31200** | Coniferous tree cover | 70, 90, 91, 92, 100, 101, 32 |

| | | |
|---|---|---|
| **31202** | Coniferous tree cover 30-60% | 70, 90, 91, 92, 100, 101, 32 |
| **31203** | Coniferous tree cover 60-100% | 70, 90, 91, 92, 100, 101, 32 |
| **31300** | Mixed tree cover | 40, 41, 50, 60, 100, 101, 32, 70, 90, 91, 92, 100, 101, 32 |
| **31400** | Tree cover in agricultural context | 32 |
| **31450** | Tree cover in urban context | 190 |
| **31500** | Green linear elements - linear woody features | 130,131,132,133,134,136 |
| **31600** | Patchy woody features | 130,131,132,133,134,136 |
| **31610** | Additional woody features | 130,131,132,133,134,136 |
| **32000** | Scrub and shrubland | 130,131,132,133,134,136,152 , 110,120 |
| **32100** | Alpine and sub-alpine natural grassland | 140,141,144,151,110,120 |
| **32200** | Moors and Heathland - other scrubland | 130,131,132,133,134,136,152 , 110,120 |
| **32300** | Sclerophyllous vegetation | 130,131,132,133,134,136,152 , 110,120 |
| **33100** | Beaches, dunes, sands | 200, 202,150,151,152 |
| **33200** | Bare rocks and rock debris | 200, 202,150,151,152 |
| **33300** | Sparsely vegetated land | 150,151,152 |
| **33500** | Permanent snow covered surfaces | 220 |
| **41000** | Wetland (permanent wet areas) - inland marshes | 180,185 |
| **41200** | Peatbogs | 180,185 |
| **42100** | Coastal salt marshes | 180,185 |
| **42200** | Intertidal flats | 180,185 |
| **51000** | Water bodies | 210 |
| **51100** | Rivernetwork | 210 |
| **51200** | Riverbed > 10m width | 210 |
| **52100** | Lagoons and Estuaries | 210 |

Functional landscape connectivity modeling (Omniscape):

For a subset of 62 species with dispersal distances within the landscape extent, we modeled functional landscape connectivity using the Omniscape algorithm (McRae et al., 2016), which is based on circuit theory. For each species, two key inputs were generated from the 5m downscaled habitat suitability map:

- Source map: Pixels with a habitat suitability probability exceeding a species-specific optimal threshold were defined as "sources" from which movement could originate.

- Resistance map: The habitat suitability map was converted into a resistance surface using a non-linear transformation function (Keeley et al., 2016), where higher suitability corresponds to lower resistance to movement.

Omniscape then calculates the cumulative current flow across the landscape, using a moving window radius defined by the species' mean dispersal distance. This results in a map of "cumulative current," which represents the probability of landscape use by a moving organism. For the remaining 164 species with very broad dispersal capacities, landscape connectivity was assumed to be complete at the scale of analysis, and their downscaled habitat suitability map was used directly in the final step. The final output for all species is a connectivity-corrected habitat suitability map at 5m resolution.

##### S3.3. Pollination model

Pollination potential was assessed as the connectivity of the landscape for a pollinator archetype, i.e. defined by characteristics broadly representing pollinators. Specifically, habitat suitability was defined as 0, 0.5 or 1 depending on the suitability of the habitat (non-habitat, partial or full habitat), based on Schulp et al. (2014) (Table S7). While forests are typically not considered as habitat, forest edges (within 10m of a border to agricultural or grassland areas) were considered as partial habitat (Schulp et al. 2014).

**Table S 7.** Suitability habitat for pollinators and demand for pollination scores used for the pollination model.

| CODE | DESC | Pollinator | Demand of pollination |
|---|---|---|---|
| **21222** | Sugar beet | 0 | 1 |
| **21231** | Sunflower | 0 | 1 |
| **21232** | Rape and turnip rape | 0 | 1 |
| **21233** | Soya | 0 | 1 |
| **21240** | Dry pulses | 0 | 1 |
| **21250** | Fodder crops (cereals and leguminous) | 0 | 1 |
| **21290** | Bare arable land | 0 | 0 |
| **22000** | Permanent crops | 0 | 1 |
| **22100** | Vinyard | 0 | 1 |
| **22200** | Orchard | 0.5 | 1 |
| **23100** | Managed grassland - Pastures | 0.5 | 0.5 |
| **23200** | Seminatural grassland - Meadows | 0.5 | 0.5 |
| **31100** | Broadleaf tree cover | 0.5 | 0.5 |
| **31102** | Broadleaf tree cover 30-60% | 0.5 | 0.5 |
| **31103** | Broadleaf tree cover 60-100% | 0.5 | 0.5 |
| **31200** | Coniferous tree cover | 0.5 | 0.5 |
| **31202** | Coniferous tree cover 30-60% | 0.5 | 0.5 |
| **31203** | Coniferous tree cover 60-100% | 0.5 | 0.5 |

| | | | |
|---|---|---|---|
| **31300** | Mixed tree cover | 0.5 | 0.5 |
| **32000** | Scrub and shrubland | 1 | 0.5 |
| **32100** | Alpine and sub-alpine natural grassland | 0.5 | 0.5 |
| **32200** | Moors and heathland - other scrubland | 1 | 0.5 |
| **32300** | Sclerophyllous vegetation | 1 | 0.5 |

The habitat suitability map was then converted to a resistance map based on distance to partial or full habitats using an exponential decay function weighted by habitat quality (partial or full). The resistance was computed as follows:

$$R = 100 - 99 * \max(0.5 * e^{-r\,d_1}, e^{-r\,d_2})$$

The decay rate r was set such that the decay function reached half probability at the median foraging distance $D_{median}$ in km:

$$0.5 = e^{-r\,D_{median}} \text{ therefore } r = \frac{\ln 2}{D_{median}} \text{ setting } D_{median} = 0.5 \text{ yields } r \approx 1.4$$

R is the resistance, d1 and d2 are distances (in km) to the closest pixel of partial and full habitat, respectively (Figure S3). This means pixels of non-habitat became more resistant the further they were located from habitat pixels. Finally, the passage probability of pollinators in each pixel was calculated using the Omniscape software, using a moving windows radius of 500m (i.e., assumed to be the average dispersal distance).

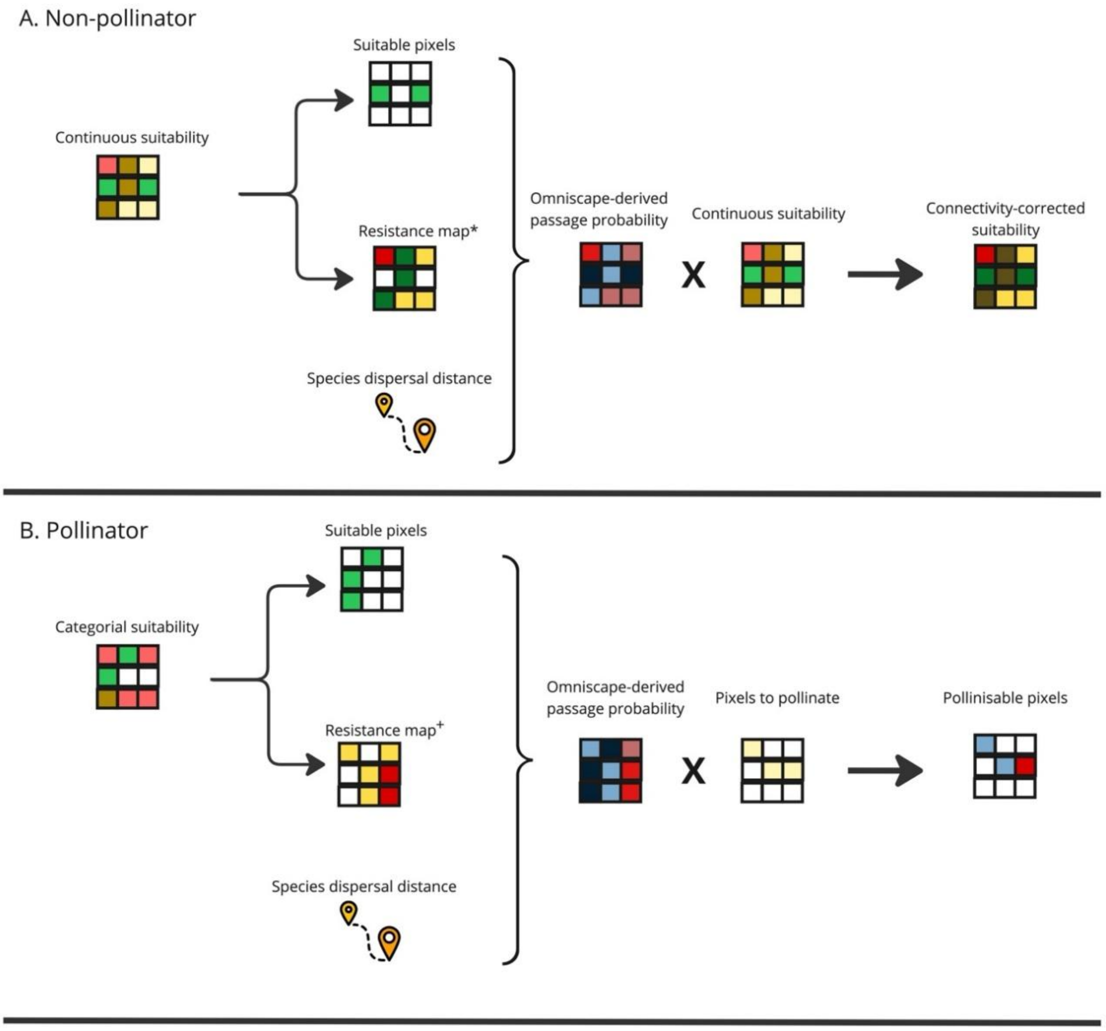

**Figure S 3.** Pollinator connectivity rationale. *derived from R = 100 - 99* (1- exp (-4*S))/(1- exp(-4), R is resistance, S is continuous suitability. + derived from R = 100 - 99*max(0.5*exp(-1.4*d1), exp(-1.4*d2)), R is resistance, d1 and d2 are distances (in km) to closest pixel of habitat type 1 and 2, respectively; assuming median foraging distance is of 500meters. Here, foraging and dispersal distances are both equal to 500meters.

To restrict this pollination potential to where it is demanded, we also classified all land uses as 0, 0.5 or 1 depending on their independence, partial or full dependence on pollination (Schulp et al. 2014, Klein et al. 2007). The final ecosystem service map was then calculated by multiplying the passage probability by the demand.

##### S3.4. Mosquito control model

This model is based on averaging suitability-weighted connectivity (or suitability alone) maps of a group of 81 species potentially linked to mosquito control (see Table S5 for species). To update the model and incorporate the demand and access of this ecosystem service, we used mosquito observation points for *Aedes albopictus* and *Culex pipiens*. These two species were selected because they are the most directly associated with potential human nuisance in the region. While other species may also cause nuisance, they are more commonly related to animals than to humans.

We contacted the Interdepartmental Mosquito Control Authority in Grenoble to obtain data on the presence of these two species. This institution is responsible for monitoring mosquito populations and notifying potential health risks linked to mosquitoes. We obtained two datasets: one for the presence of *Aedes albopictus*, which included an Excel file with 22,958 recorded observation points, and another for *Culex pipiens*, with 3,204 recorded points.

After clipping the data to fit the study area, we retained a total of 4,318 points for *Aedes albopictus* and 892 for *Culex pipiens*. We merged both datasets and applied a habitat modelling approach using Biomod2 (Thuiller et al., 2016) following the methodology in (Sherpa et al., 2020) to create a raster layer representing the habitat suitability for both species as a measure of demand for mosquito control.

Additionally, we created a 1 km buffer around urban areas, recognizing that human mobility is a key factor in mosquito exposure. This buffer distance was informed by a participatory mapping approach conducted in Grenoble, where over 600 participants were asked about various ecosystem-related benefits, including their willingness to travel to nearby ecosystems for regular recreational activities. To create the buffer, we used the land use-land cover layer and selected the codes 11100, 11200, 11300, 11400, 12100 corresponding with areas where people live or work. Finally, we multiplied the resulting layer by the connectivity map for mosquito control species to obtain the final mosquito control map.

##### S3.5. Hunting value model

The provision of this ecosystem service was modeled based on the aggregated, connectivity-corrected habitat suitability of 24 game species (see Table S5 for species list).

To ensure the final map reflected where hunting is a relevant activity, we integrated a demand layer based on data that considers both regulatory restrictions and other criteria affecting the potential presence of hunters. This data was provided by the regional hunters' association (Fédération Départementale des Chasseurs de l'Isère) and includes spatial information on:

- Hunting reserves: zones identified as important for the reproduction of game species, where hunting is often restricted.
- Prohibited areas: zones where hunting is explicitly forbidden.
- A 150-meter regulatory buffer around buildings, where hunting is generally not allowed.

As these regulations can be nuanced (e.g., hunting may be permitted under certain conditions in reserves for population control), we incorporated these criteria by applying the following weighting adjustments to the aggregated species connectivity map:

- Areas classified as urban or artificial in the land use map were assigned a value of 0.
- Areas where hunting is explicitly prohibited were assigned a value of 0.1.
- Reserve areas were assigned a value of 0.2.
- Areas within the 150-meter buffer around buildings were assigned a value of 0.5.
- All other areas retained their original value from the connectivity map.

This process results in a final map that represents hunting value, spatially filtered by the social and regulatory context that governs the activity.

##### S3.6. Biological control model

The provision of this ecosystem service was modeled based on the aggregated, connectivity-corrected habitat suitability of 78 species known to prey on agricultural and garden pests (see Table S5 for species list). We integrated demand by considering that this ecosystem service is primarily relevant in areas where pests have a direct impact on human activities: agricultural lands and urban green spaces (e.g., parks and private gardens).

To implement this, the aggregated species connectivity map was multiplied by a demand mask that restricted the ecosystem service to these specific land-cover classes. All areas outside of these designated crop and urban green space areas were masked to 0. The land-use codes from the EUSALP map used to create this mask were: 14100, 21211, 21212, 21213, 21214, 21215, 21216, 21218, 21221, 21222, 21230, 21231, 21232, 21233, 21240, 21250, 21290, 22100, 22200, 23100, 23200, and 32100. These codes correspond to the land use and land cover categories used in the food production model, with the addition of urban areas.

##### S3.7. Seed dispersal model

Seed dispersal is a fundamental ecological process that underpins the resilience and regeneration of plant communities, contributing to the long-term maintenance of habitat structure across the entire landscape. The provision of this ecosystem service was modeled based on the aggregated, connectivity-corrected habitat suitability of 81 frugivorous species (birds and mammals) known to disperse native plant seeds (see Table S5 for species list).

For this particular ecosystem service, demand was considered to be ubiquitous and non-localized. Unlike ecosystem services targeted at specific human activities (like pollination for crops), seed dispersal provides a foundational benefit to all natural and semi-natural ecosystems. Its importance is not restricted to specific locations but is distributed across the landscape, wherever ecosystem regeneration and resilience are required. Consequently, no spatial demand mask was applied, and the aggregated species connectivity map was used directly as the final output for this ecosystem service.

##### S3.8. Emblematic species model

This ecosystem service represents the cultural and existence value derived from the presence of iconic and charismatic species, which are central to regional identity and nature-based tourism.

The provision of this ecosystem service was modeled based on the aggregated, connectivity-corrected habitat suitability of 96 species considered emblematic in the region (see Table S5 for species list). Species were selected as emblematic based on a combination of criteria, including their conservation status (e.g., species listed in the EU Habitats and Birds Directives), their importance for nature-based tourism (e.g., frequent mentions on regional tourism websites), and high public interest as indicated by citizen science observation rates (O'Connor et al., 2025).

Similar to seed dispersal, the demand for this ecosystem service was considered ubiquitous and non-localized. The value people derive from the existence of emblematic species is not confined to the specific locations where they might be observed, but extends across the entire landscape as part of the region's natural heritage. Consequently, no spatial demand mask was applied, and the aggregated species connectivity map was used directly as the final output.

#### S4. Methodological details: Structural buffers and sampling protocol

##### S4.1. Structural buffers creation

The use of structural buffers allows for a more ecologically and socio-economically meaningful analysis of gradients compared to conventional Euclidean distance buffers. These buffers define zones based on a "cost of travel" through the landscape structure, ensuring that all points within a given buffer zone are, in a practical sense, "equidistant" from the protected area border in terms of accessibility and landscape permeability (See Figure S4A).

The entire cost-distance analysis was performed iteratively using Python libraries including NumPy, GDAL, scikit-image, and SciPy. The process was conducted twice to assess these structural gradients both outwards from and inwards towards protected areas.

- Generation of the structural resistance surface: In the first iteration, a raster representing landscape resistance was configured to assign a value of zero within protected area boundaries, designating them as the least-cost origins for movement outwards. The resistance value for pixels outside the protected areas was calculated as a composite of land cover and topographic factors using the formula: Final Resistance = 2 * (Land-Cover Resistance) + Topographic Resistance. Topographic resistance was derived from a 5m DEM and classified into three categories based on slope (<50%, 50-100%, and >100%). Land-cover resistance values are detailed in Table S8.
- Cumulative cost raster: Following the creation of the resistance raster, the Geometric Minimum Cost Path algorithm from scikit-image (skimage.graph.mcp) was executed to generate a cumulative cost raster representing the structural resistance gradient emanating from the protected areas.
- Buffer delineation: This cumulative cost raster was then used to delineate five structural buffers both outwards (positive buffer numbers) and inwards (negative buffer numbers, derived from a complementary second iteration). The zones were defined based on the following cumulative resistance thresholds: Buffer 1/-1 (resistance ≤ 100); Buffer 2/-2 (101-200); Buffer 3/-3 (201-400); Buffer 4/-4 (401-800); and Buffer 5/-5 (>800). These thresholds define zones of progressively increasing landscape isolation relative to the protected area.

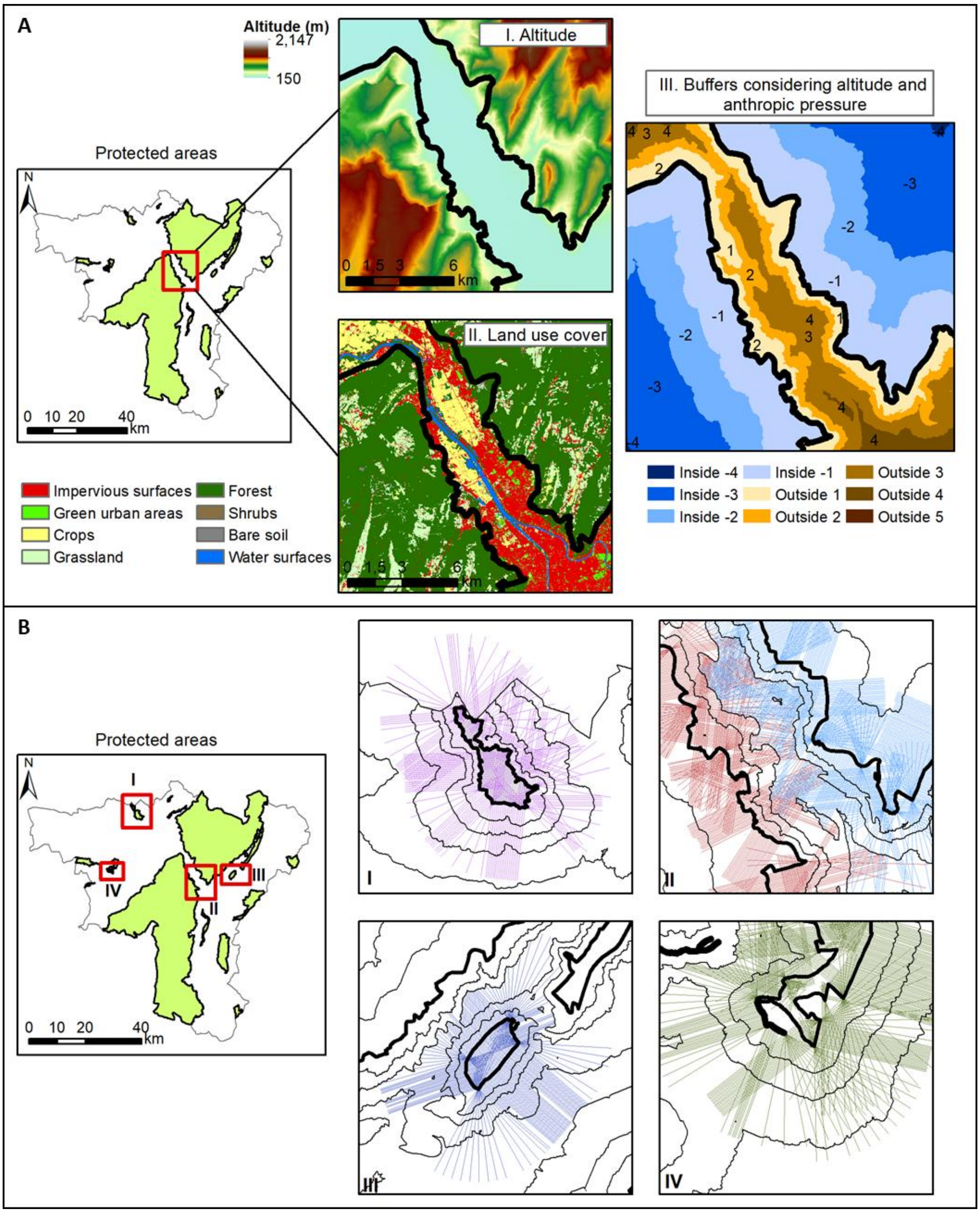


**Figure S 4.** Illustration of the methodology for the generation of structural buffers and the transect sampling procedure. Panel (A) illustrates the process for creating structural buffers. The method integrates two main sources of landscape resistance based on the structure of the landscape: (I) topographic resistance, derived from an altitude map where steep slopes impose a higher cost to movement; and (II) resistance associated with land use/land cover, where each category is assigned a specific resistance value (refer to Table S8 for details). The final map (III) displays the resulting cumulative resistance buffers for an example of two closely located protected areas (thick black lines). Buffers are generated both inwards (shades of blue) and outwards (shades of orange and brown) from the border, and their variable width reflects the heterogeneous landscape structure and resistance. Panel (B) then illustrates the logic for transect generation and filtering across four representative scenarios, where thick black lines denote the protected area borders and thin lines denote the structural buffers. For an isolated protected area (I), transects can reach their maximum length without being truncated, as there is no interference from neighboring protected areas. In contrast, the detailed view in (II) shows how the effect of overlapping buffers truncates the transects; the blue lines originating from the right protected area stop before touching the buffers of the left protected area, ensuring that each transect exclusively analyzes the gradient from its source protected area. The methodology also accounts for complex geometries, such as a protected area with sharp vertices (III), where multiple transects are generated to ensure complete border coverage while still applying filtering rules against nearby areas. Finally, for protected areas with winding, serpentine shapes (IV), transects are generated from a central line towards the border to prevent them from self-intersecting or re-entering their own protected area, thus guaranteeing the validity of the measured gradient.

**Table S 8.** Resistances assigned to each land use/land cover class to create the resistance layer for structural buffer calculations.

| CODE | DESC | Cost |
|---|---|---|
| **11000** | Artificial surfaces and constructions | 1 |
| **11100** | Dense settlement area | 1 |
| **11200** | Low density settlement area | 0.9 |
| **11300** | Builtup area | 1 |
| **11400** | Open settlement area | 0.9 |
| **12100** | Industrial and commercial zones | 1 |
| **12210** | Roads motroways and trunks | 1 |
| **12220** | Road networks | 1 |
| **12221** | Roads tertiary and others | 1 |
| **12230** | Railways train tracks | 0.9 |
| **12240** | Unpaved roads and tracks | 0.8 |
| **14100** | Green urban areas | 0.7 |
| **21000** | Cultivated areas - Arable land - Annual crops | 0.5 |
| **21211** | Common wheat | 0.5 |
| **21212** | Durum wheat | 0.5 |
| **21213** | Barley | 0.5 |
| **21214** | Rye | 0.5 |
| **21215** | Oats | 0.5 |
| **21216** | Maize | 0.5 |
| **21218** | Triticale | 0.5 |
| **21221** | Potatoes | 0.5 |
| **21222** | Sugar beet | 0.5 |
| **21230** | Other non permanent industrial crops | 0.5 |
| **21231** | Sunflower | 0.5 |
| **21232** | Rape and turnip rape | 0.5 |

| | | |
|---|---|---|
| **21233** | Soya | 0.5 |
| **21240** | Dry pulses | 0.5 |
| **21250** | Fodder crops (cereals and leguminous) | 0.5 |
| **21290** | Bare arable land | 0.5 |
| **22000** | Permanent crops | 0.5 |
| **22100** | Vinyard | 0.5 |
| **22200** | Orchard | 0.5 |
| **23100** | Managed grassland - Pastures | 0.4 |
| **23200** | Seminatural grassland - Meadows | 0.4 |
| **31100** | Broadleaf tree cover | 0.1 |
| **31102** | Broadleaf tree cover 30-60% | 0.1 |
| **31103** | Broadleaf tree cover 60-100% | 0.1 |
| **31200** | Coniferous tree cover | 0.1 |
| **31202** | Coniferous tree cover 30-60% | 0.1 |
| **31203** | Coniferous tree cover 60-100% | 0.1 |
| **31300** | Mixed tree cover | 0.1 |
| **31400** | Tree cover in agricultural context | 0.3 |
| **31450** | Tree cover in urban context | 0.8 |
| **31500** | Green linear elements - linear woody features | 0.2 |
| **31600** | Patchy woody features | 0.2 |
| **31610** | Additional woody features | 0.2 |
| **32000** | Scrub and shrubland | 0.1 |
| **32100** | Alpine and sub-alpine natural grassland | 0.1 |
| **32200** | Moors and heathland - other scrubland | 0.1 |
| **32300** | Sclerophyllous vegetation | 0.1 |
| **33100** | Beaches, dunes, sands | 0.1 |
| **33200** | Bare rocks and rock debris | 0.2 |
| **33300** | Sparsely vegetated land | 0.2 |
| **33500** | Permanent snow covered surfaces | 0.4 |
| **41000** | Wetland (permanent wet areas) - inland marshes | 0.1 |
| **51000** | Water bodies | 0.3 |
| **51100** | Rivernetwork | 0.3 |

| | | |
|---|---|---|
| **51200** | Riverbed > 10m width | 0.2 |

##### S4.2. Sampling methods details

The process began with the simplification of the protected area polygons using the Douglas-Peucker algorithm (buffer radius = 100m, simplification tolerance = 300) to facilitate the subsequent geometric operations.

Following simplification, the border of each polygon was divided into a set number of equal-length segments. This border segmentation ensures a consistent 360-degree representation regardless of the protected area's shape. To maintain a comparable level of thematic detail across the study, the number of segments was scaled to the protected area's size: 20 segments for large (>10,000 ha), 10 for medium (1,000-10,000 ha), 5 for small (100-1,000 ha), and 3 for very small (<100 ha) protected areas.

Once the segments were defined, we proceeded with transect generation. Along the length of each segment, perpendicular transect lines were generated at 25m intervals, extending a maximum of 4 km both inwards and outwards from the border.

To ensure the spatial validity and coherence of these sampling lines, all generated transects were then subjected to a rigorous automated filtering process. This crucial step, implemented using the GeoPandas and Shapely libraries in Python, ensures that each transect accurately represents a gradient away from a single, specific border segment. The filtering was governed by the following rules (see Figure S4B):

- Polygon intersection check: Transects that incorrectly crossed back into their own protected area polygon interior after exiting were discarded.
- Minimum length threshold: Transects shorter than a 20m minimum length after adjustment were discarded as too short to capture a meaningful border gradient.
- Buffer zone overlap avoidance: To prevent erroneous sampling of gradients influenced by neighboring protected areas, transects were checked for overlaps with the structural buffers of other PAs. If a transect intersected a buffer zone from another PA, the line was truncated at that point of second intersection.
- Polygon overlap discard: Transects crossing the protected area polygon more than once (indicating geometric anomalies in line generation) were discarded.

Each transect that passed this filtering process was then assigned to its corresponding border segment, ready for the extraction of bundle values and the subsequent gradient analysis.

##### S4.3. Border typology generation: Algorithmic details

The classification of each segment–bundle profile into the A-E typology was performed using an automated method based on polynomial regression.

Profile construction: For each border segment and ecosystem service bundle, a single profile was constructed from multiple data points. Each data point represents the mean normalized ecosystem service value calculated across all transects within that segment for a specific structural buffer zone. The x-axis for the regression represents the ordinal buffer level, from the innermost buffer inside the protected area to the outermost buffer outside.

Polynomial regression: For each of these profiles, we performed polynomial regressions to mathematically characterize its shape.

- Response and predictor variables: The response variable was the mean normalized ecosystem service bundle value. The predictor variable was the buffer level, treated as an ordinal variable.

- Model fitting: We fitted both first-degree (linear) and second-degree (quadratic) polynomial models. The best-fitting model was adaptively selected for each profile based on which model yielded the lower Mean Squared Error.

Group assignment based on curve shape: Once the best-fitting model was selected, each profile was classified into one of five types based on a set of rules applied to the model's coefficients. All algorithmic parameters and implementation specifics are available in the code repository.

- Types A, B, and C (linear trends): If a first-degree polynomial provided a good fit (Mean Squared Error < 0.05), the profile was classified as a linear trend based on its slope:
  - Type A ('Flat Gradient'): Slope coefficient absolute value was below a flatness threshold of 0.01.
  - Type B ('Increasing Gradient'): Slope was positive (≥ 0.01).
  - Type C ('Decreasing Gradient'): Slope was negative (≤ -0.01).
- Types D and E (quadratic trends): If a second-degree polynomial was selected, the profile was classified as a quadratic trend based on its concavity (the sign of the quadratic term):
  - Type D ('Boundary Depression'): Concave-up.
  - Type E ('Interface Peak'): Concave-down.

**S5. Methodological details: Statistical analysis of landscape drivers**

To analyze the effect of crossing the protected area boundary on landscape metrics, we employed a two-level statistical strategy. This approach combines the power of Linear Mixed Models for global testing with the specificity of paired post-hoc tests for detailed comparisons.

- Landscape metrics calculated: For each border segment, we calculated a suite of landscape metrics for the aggregated areas "inside" and "outside" the protected area.
  - Composition: We first aggregated the detailed EUSALP land-use codes into three functional macro-categories: natural (e.g., forests, natural grasslands, wetlands), semi-natural (primarily agricultural lands), and artificial (e.g., urban areas, infrastructure). We then calculated the proportion of the area covered by each of these macro-categories.
  - Structure/Diversity: We calculated Shannon diversity and Pielou's evenness at two levels: using the three macro-categories and using the full, detailed land use code classification. We also calculated Dominance as the proportion of the most frequent land use type.
  - Configuration/Fragmentation: We measured spatial autocorrelation using the mean Moran's I value across all land cover codes. This serves as a robust proxy for fragmentation, where higher values indicate a more continuous, less fragmented landscape.
- Level 1: Global test with linear mixed models: The primary analysis used Linear Mixed Models to determine if the pattern of change in a given metric from "inside" to "outside" (Location) was consistently different across the five border types (Border Type).
  - Model structure: The core model tested the significance of the interaction term: Landscape Metric ~ Location * Border Type + (1 | Protected_Area / Border Segment).
  - Random effects: We included Protected Area and Border Segment as nested random effects to account for the non-independence of measurements.

  - Model validation: The assumptions of normality and homoscedasticity of the model residuals were visually inspected for each model. A significant interaction term ($p < 0.05$) justified proceeding to post-hoc tests.
- Level 2: Post-hoc paired comparisons: When the interaction term in the Linear Mixed Model was significant, we performed paired tests to identify which specific border types showed a significant difference between their "inside" and "outside" landscape characteristics.
  - Test selection: For each comparison, we calculated the paired differences (value outside – value inside). A Shapiro-Wilk test was then applied to check the normality of these differences. In all cases, the differences were found to be non-normally distributed ($p < 0.05$).
  - Final test: Consequently, we consistently used the Wilcoxon signed-rank test, the appropriate non-parametric alternative to a paired t-test, to ensure the robustness of the specific findings reported in the Results section.

**S6. Methodological details: Robustness analysis**

To evaluate the internal consistency of the ecosystem services bundles and ensure that the aggregated results accurately represent the underlying individual dynamics, we conducted a validation analysis across all border segments.

**S6.1. Geometric similarity analysis (correlation)**

We measured the shape similarity between the gradient curves of individual ecosystem services and their corresponding management bundles. Because the gradients are represented by polynomial equations, it was necessary to generate predicted values at specific intervals to perform a quantitative comparison. We reconstructed each equation over a standardized range from -4 to 5, calculating the predicted values at ten discrete points corresponding to the structural buffer levels. This process allowed us to generate a series of numerical data pairs for each individual ecosystem service and its corresponding bundle, which were then used to calculate the Pearson correlation coefficient (r). This metric reveals the extent to which the shape of an individual ecosystem service gradient mirrors (r near 1), moves independently of (r near 0), or opposes (r near -1) the average trend of its corresponding bundle.

**S6.2. Categorical similarity analysis (typology agreement)**

We assessed the consistency of the final classification to determine how often an individual ecosystem service was assigned the same gradient typology (Types A–E) as its corresponding bundle. For every border segment and individual ecosystem service, we calculated a categorical similarity score by identifying matches between the service-level typology and the bundle-level typology. This percentage of agreement indicates whether the aggregated bundle is a representative indicator of its constituent individual ecosystem services or if specific ecosystem services exhibit idiosyncratic patterns that diverge from the group average.

**6.3. Place-based ecosystem services analysis**

We conducted a sensitivity analysis to test the influence of different ecosystem service types on our results, particularly to address the role of static ecosystem services. We focused on the rural bundle, as it is the only one containing the two ecosystem services with solely local (pixel-based) determinism: food production (high in the agricultural matrix outside protected areas) and carbon storage (high in natural habitats inside).

To do this, we created a new, alternative rural bundle that included only the three connectivity-based ecosystem services (pollination, biological control, hunting value) and re-ran our entire border classification.

#### S7. Supporting information for results interpretation

##### S7.1. Additional information for ecosystem services interpretation

To facilitate detailed border analysis and management strategy adaptation for specific ecosystem services, our code automatically generates visualizations for each protected area and ecosystem service. Figure S5 provides an illustrative example for Protected Area 12, Border Segment 3. However, the code (https://doi.org/10.5281/zenodo.17876914) produces one figure per segment per protected area for comprehensive analysis.

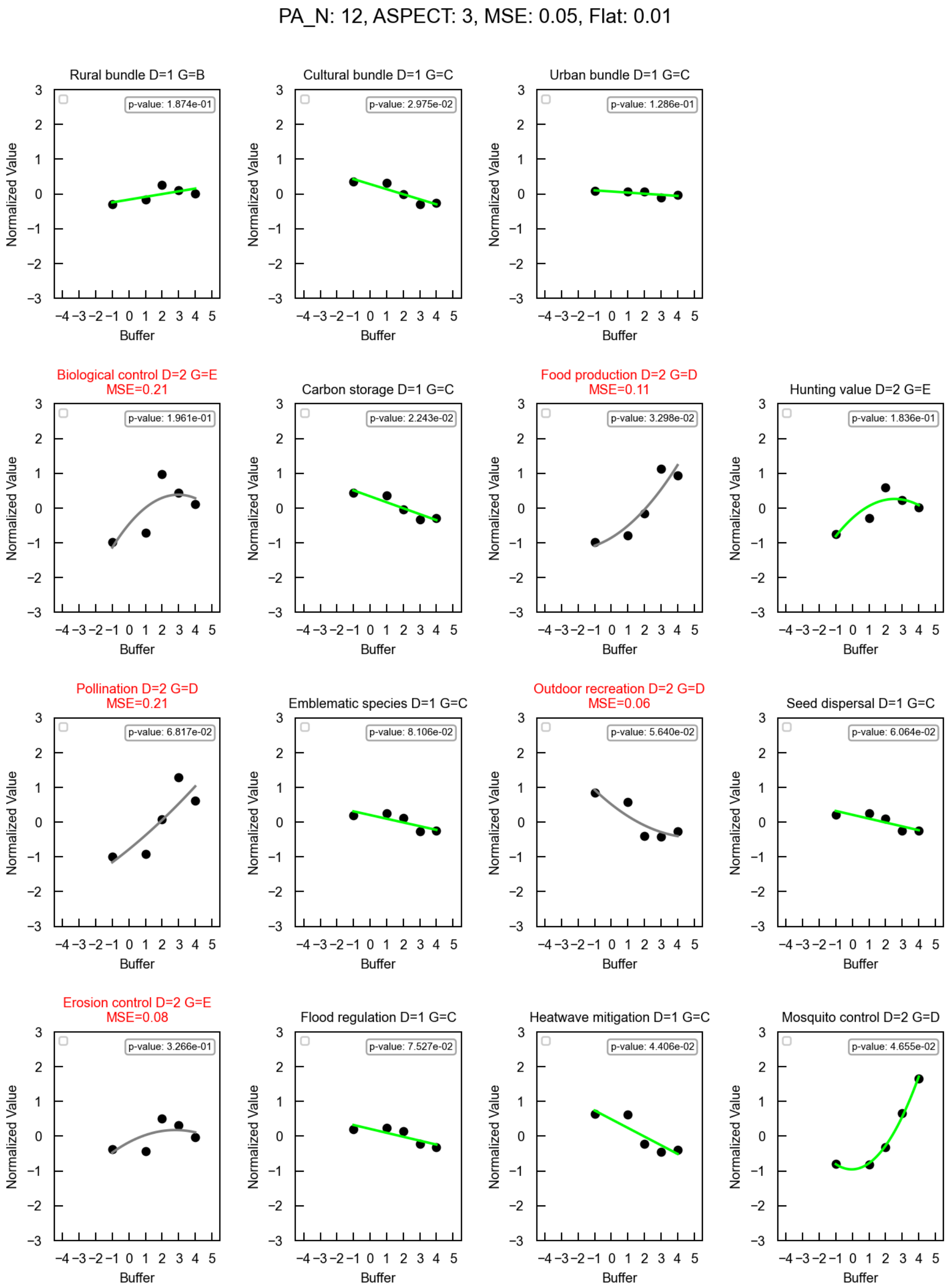


**Figure S 5.** Example of a figure generated using our methodology for the detailed interpretation of borders and ecosystem services. The figure represents an example for Protected Area 12 (PA_N), border segment 3 (Aspect), and a flat rate of 0.01 (which defines the slope). Each graph displays the polynomial degree (D) and the type (G). The mean square error (MSE) is shown when it exceeds the defined threshold (0.05).

The Y-axis represents the normalized value. The points defining the lines correspond to the average value of each line crossing the border segment intersected by the buffers. Green lines indicate a good fit based on the defined criteria, while gray lines represent fits where the mean square error exceeds the 0.05 threshold.

Figure S6 shows the results of the ecosystem service maps without integrating demand and access criteria and protected area perimeters to support the interpretation of the results for the 12 modelled ecosystem services.

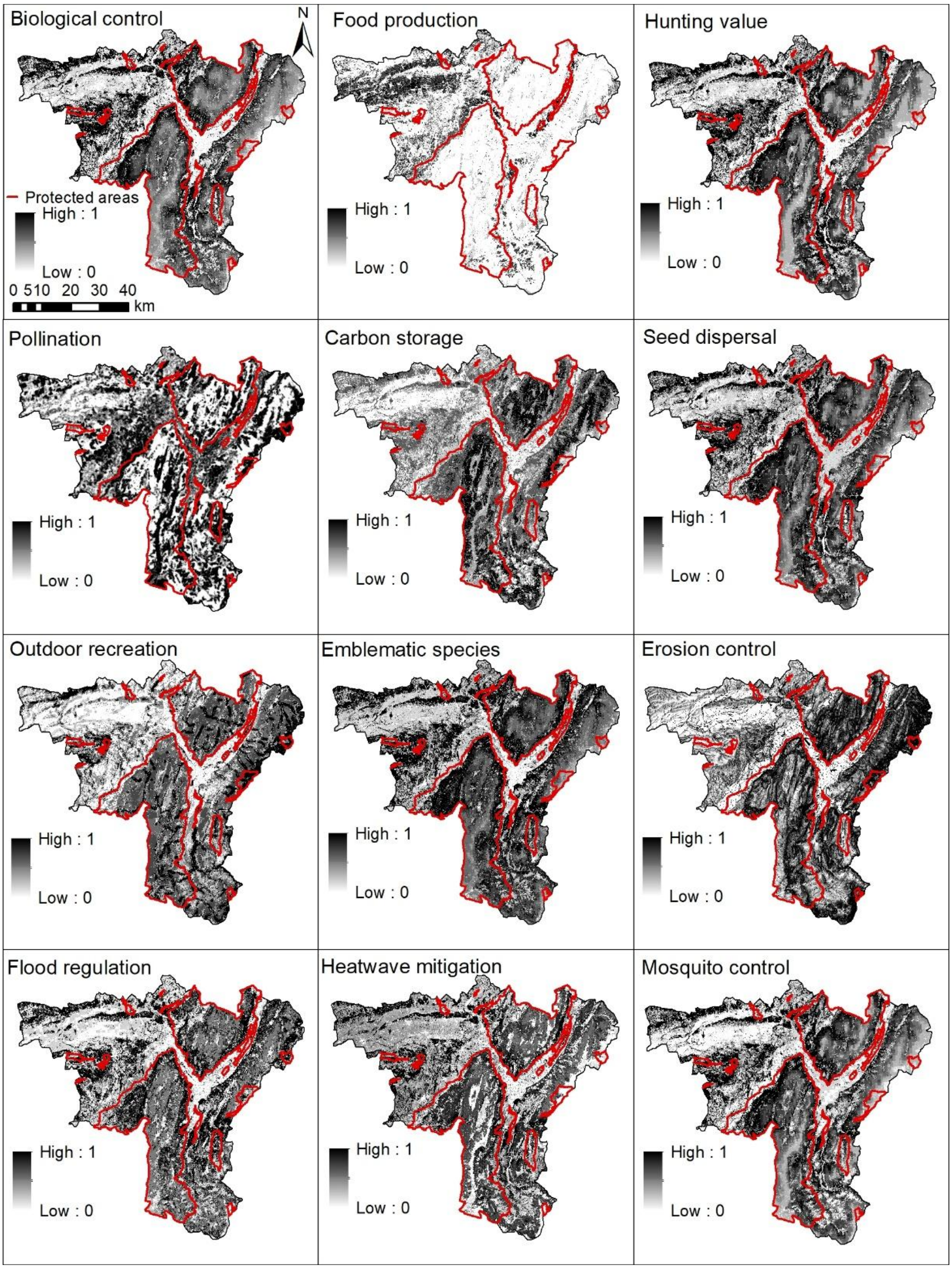


**Figure S 6.** Results of ecosystem services models without integrating demand and access variables. These maps represent the potential value of the ecosystem services analysed. Details on methods are provided in Appendix S3.

Figure S7 shows the results of the ecosystem service maps including access and demand and protected area perimeters to support the interpretation of the results for the 12 modelled ecosystem services.

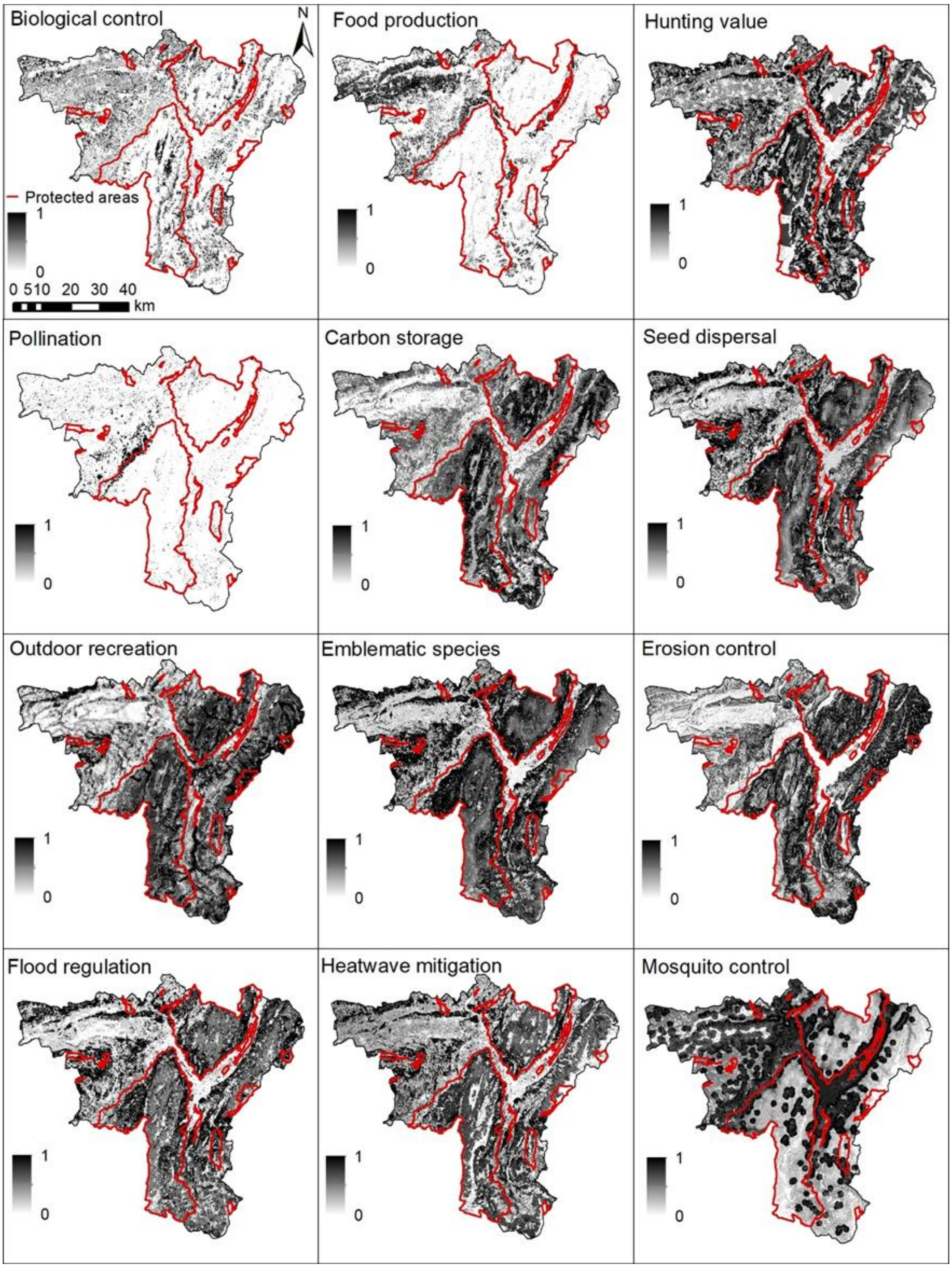


**Figure S 7.** Results of the final ecosystem service models, including demand and access criteria. The rural bundle includes biological control, food production, hunting value, pollination, and carbon storage. The cultural bundle includes seed dispersal, outdoor recreation, and emblematic species. The urban bundle includes erosion control, flood regulation, heatwave mitigation, and mosquito control. The maps reflect the specific variables related to the supply, demand, and access of ecosystem services, as explained in each model section of Supporting Information-E. Food production, carbon storage, seed dispersal, emblematic species, erosion control, flood regulation, and heatwave mitigation do not show changes compared to Figure S6.

In addition to the interpretation of each border type, we developed an additional figure to visually assess ecosystem services values and bundle values around protected areas and borders. Figure S8 shows the distribution of pixel values per bundle and ecosystem services in relation to the border segment, exemplified for the “Étangs, landes, vallons tourbeux humides et ruisseaux à écrevisses de Chambaran” protected area.

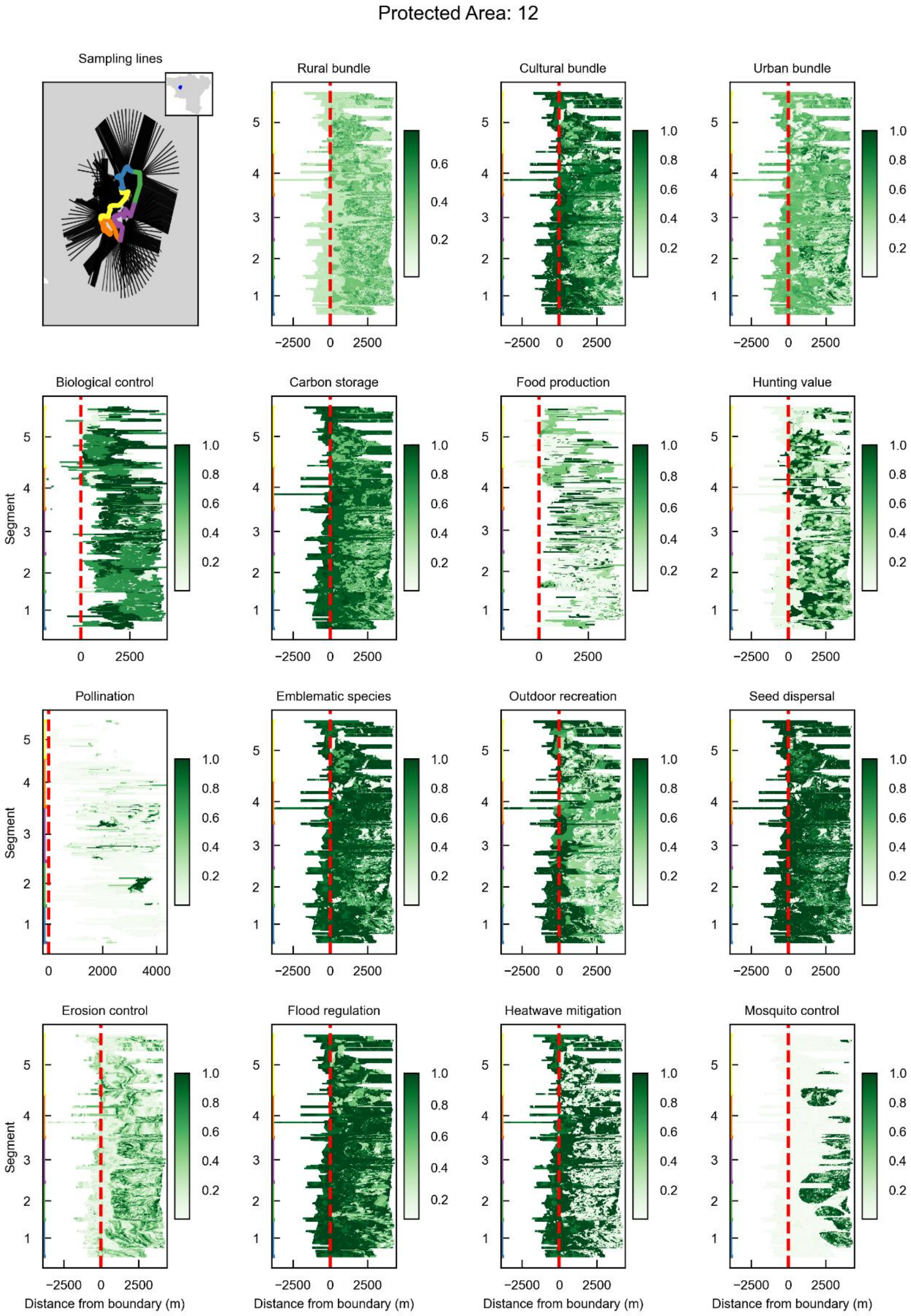


**Figure S 8.** Example Ecosystem Service and Bundle Value Distributions Around the “Étangs, landes, vallons tourbeux humides et ruisseaux à écrevisses de Chambaran” Protected Area (12). These figures are automatically generated by our methodology to provide a visual interpretation of ecosystem service and bundle value distributions outside and inside protected areas (protected area border is delineated by the red dashed line). The top-left panel displays the border segments and transect lines for the protected area. The remaining panels show heatmaps illustrating ecosystem service or bundle values (Y-axis: border segments, colours correspond with colours in the top left panel) as a function of distance from the protected area border (X-axis).

Figure S9 shows the spatial distribution of the main border type combinations presented in Figure 4 of the main manuscript.

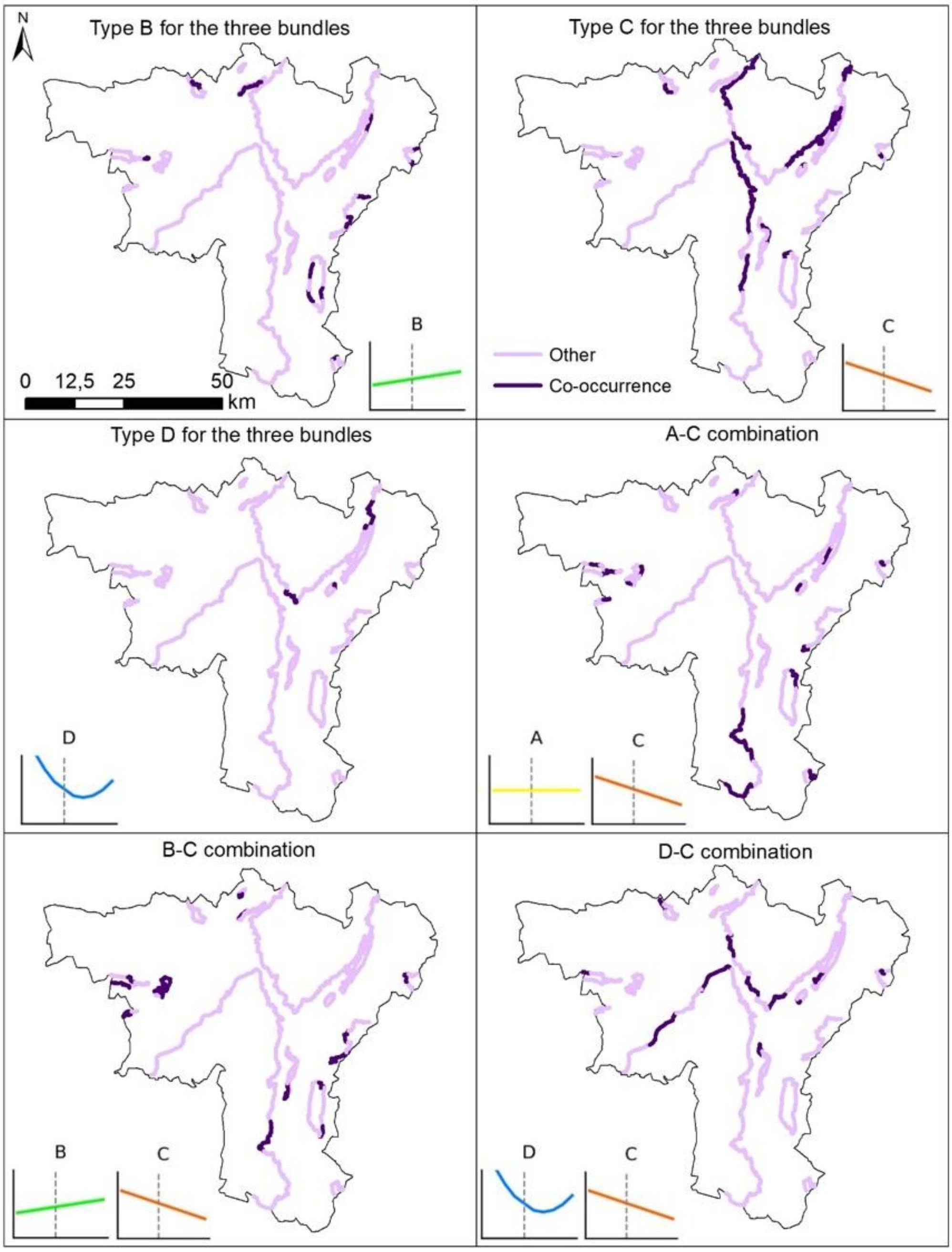


**Figure S 9.** Spatial distribution of border combinations. The titles in each subplot describe the combination being mapped in each case. These include the most frequent cases (Figure 4c in the main text) of borders with only type B, only type C, or only type D, as well as borders with combinations between A and C, B and C, and D and C. Dark purple indicates borders where the mapped interaction occurs, while light purple represents borders where no co-occurrence was detected for the mapped interaction.

##### S7.2. Robustness analysis results

The sensitivity analysis shows the degree of consistency and representativeness of the ecosystem service bundles (Table S9 and Fig. S10). The cultural bundle has the highest coherence, where the group trend mirrors the shape of the individual ecosystem service gradients almost perfectly (median r = 1.00). In the urban bundle, the high agreement and correlation of flood regulation (75.96% agreement, median r = 1.00) confirm that the bundle effectively captures the primary regulating functions in this domain, even when idiosyncratic patterns such as mosquito control (median r = -0.13) are present.

For the rural bundle, the higher level of internal divergence is not a limitation of the framework, but a reflection of the socio-ecological reality at the borders. The low consistency of food production (6.73% agreement) highlights its role as a service that often moves independently of stock-based services like carbon storage. Crucially, the sensitivity analysis confirms that the bundle-level typology is a robust diagnostic indicator for the vast majority of cases. In 97.7% of the total bundle observations across all domains, the assigned gradient type (A–E) accurately represents the dominant individual trends without being compromised by hidden opposing patterns. This high representativeness validates the use of aggregated bundles as a powerful diagnostic tool for landscape planning, while the framework simultaneously retains the ability to identify specific high-diversity segments where service-level trade-offs occur.

**Table S 9.** Categorical agreement and geometric similarity between individual ecosystem services and their corresponding management bundles. Agreement (%) represents the percentage of the 104 analyzed border segments where the individual ecosystem service was assigned the same gradient typology label (A–E) as its bundle. Median Pearson r values indicate the average shape similarity between the ecosystem service and the bundle curves across all segments. Background colors in the agreement column highlight the level of consistency, ranging from low agreement (red) to high agreement (green).

| Ecosystem service | Corresponding bundle | Agreement (%) | Median Pearson r |
|---|---|---|---|
| Emblematic species | Cultural bundle | 80.77% | 1.00 |
| Seed dispersal | Cultural bundle | 79.81% | 1.00 |
| Flood regulation | Urban bundle | 75.96% | 1.00 |
| Outdoor recreation | Cultural bundle | 63.46% | 0.99 |
| Heatwave mitigation | Urban bundle | 58.65% | 0.68 |
| Carbon storage | Rural bundle | 48.08% | 0.98 |
| Erosion control | Urban bundle | 46.15% | 0.78 |
| Hunting value | Rural bundle | 42.31% | 0.63 |
| Mosquito control | Urban bundle | 31.73% | -0.13 |
| Pollination | Rural bundle | 25.00% | 0.35 |
| Biological control | Rural bundle | 20.19% | 0.03 |
| Food production | Rural bundle | 6.73% | -0.06 |

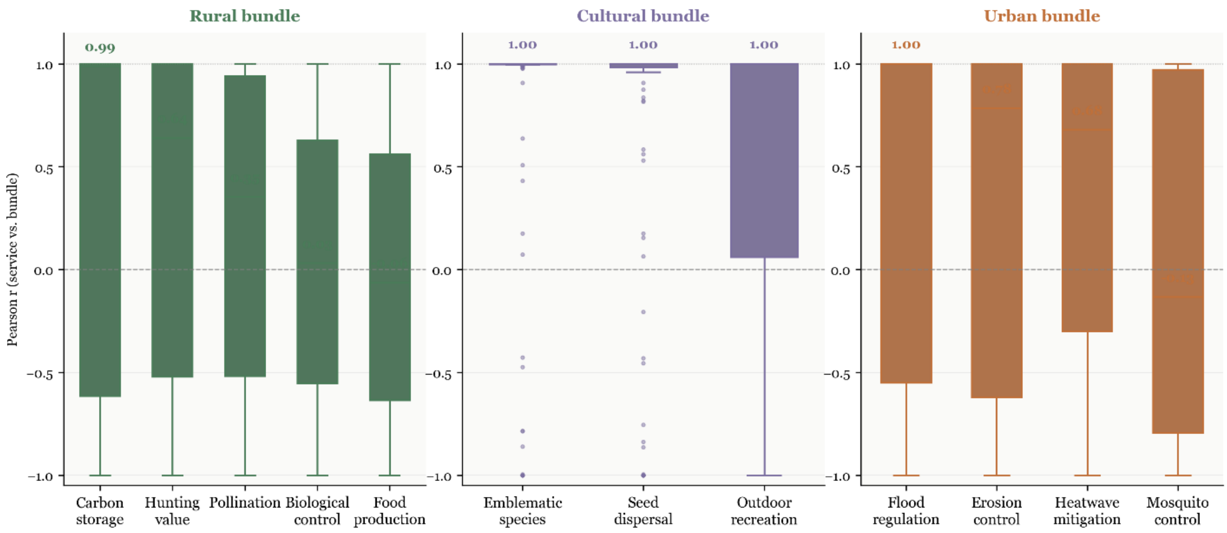


**Figure S 10.** Geometric similarity between individual ecosystem services and their corresponding management bundles. The boxplots show the distribution of Pearson correlation coefficients (r) calculated across 104 border segments. These correlations measure the similarity between the reconstructed polynomial curves of each individual ecosystem service and its corresponding bundle. Median r values are highlighted above specific boxes to identify ecosystem services that most closely mirror the bundle trend. Values near 1.00 indicate that the individual ecosystem service gradient follows the same pattern as the aggregated bundle, while values near 0 or negative values indicate independent or opposing spatial trends.

##### S7.3. Detailed statistical results for landscape drivers

This section details the statistical analysis performed to identify the landscape configurations driving the different border typologies, supporting the findings presented in the main text (Results section, "Landscape drivers of border patterns").

The analysis was conducted in two stages. First, we used Linear Mixed Models (LMM) to test for overall significant differences in landscape patterns among the border types. Second, where significant differences were found, we used post-hoc paired tests to identify the specific nature of those differences. We analyzed a suite of metrics representing three key aspects of landscape structure. Composition was measured as the proportion of three aggregated land-cover classes: natural, semi-natural (primarily agricultural), and artificial. Structural diversity was assessed using the Shannon Diversity Index (H). Finally, landscape configuration was measured using spatial autocorrelation (Moran's I) as a proxy for fragmentation, where higher values indicate a more continuous and less fragmented landscape.

For each of the nine primary analyses (three metrics for each of the three bundles), we fitted a Linear Mixed Model. The model tested if the change in a given landscape metric across the protected area boundary differed significantly among the five border types. The boundary itself was represented by a two-level factor we term Location, which distinguishes between the aggregated areas "inside" and "outside" the protected area. The key result was the significance of the interaction term (Location * Border Type). In all nine models, this interaction term was highly significant ($p < 0.001$). This confirms that the five border types (A-E) correspond to genuinely different and statistically distinct patterns of landscape transition across the border. This significant global result justifies the subsequent post-hoc analysis. Table S10 presents the results of the post-hoc Wilcoxon signed-rank tests, which detail the specific, significant differences between the "inside" and "outside" landscape characteristics for each border type.

**Table S 10.** Results of Wilcoxon signed-rank tests for landscape metrics inside vs. outside protected areas.

| Bundle | Landscape Metric | Type A | Type B | Type C | Type D | Type E |
|---|---|---|---|---|---|---|
| **Rural** | *Proportion of Natural Habitat* | < 0.05 | NS | < 0.001 | - | - |
| | *Proportion of Artificial Habitat* | NS | < 0.001 | < 0.001 | - | - |
| | *Shannon Diversity* | < 0.05 | < 0.05 | < 0.05 | - | - |
| **Cultural** | *Proportion of Natural Habitat* | NS | < 0.05 | < 0.01 | - | - |
| | *Proportion of Artificial Habitat* | NS | NS | < 0.05 | - | - |
| | *Spatial Autocorrelation (Moran's I)* | NS | NS | < 0.001 | - | - |
| **Urban** | *Proportion of Natural Habitat* | NS | < 0.05 | < 0.05 | - | - |
| | *Proportion of Seminatural Habitat* | NS | NS | < 0.05 | - | - |
| | *Spatial Autocorrelation (Moran's I)* | NS | < 0.05 | < 0.001 | - | - |

##### S7.4. The role of place-based ecosystem services in shaping border patterns

The results of the sensitivity analysis showed that excluding food production and carbon storage significantly altered the classification of the rural borders. Specifically, 35% of the segments (37 out of 104) that were originally classified as one of the three main linear types (A, B, or C) migrated to a different category. The detailed changes are summarized in Table S11. The most notable trend was a strong shift away from simple, linear profiles towards the more complex, non-linear profiles (Types D and E). For instance, a majority of the original 'Increasing Gradient' (Type B) segments became 'Interface Peak' (Type E) segments.

**Table S 11.** Migration of rural bundle border types in a sensitivity analysis excluding place-based ecosystem services. The table shows how the classification of border segments for the rural bundle changes when the two place-based ecosystem services (food production and carbon storage) are excluded from the analysis. The rows represent the original classification (using all five rural ecosystem services) for segments initially classified as Types A, B, or C. The columns show the new classification of these same segments after the exclusion**.**

| Original Type (Total N migrated) | New Classification | | | |
|---|---|---|---|---|
| | **Migrated to Type B** | **Migrated to Type C** | **Migrated to Type D** | **Migrated to Type E** |
| **Type A (N=11)** | 5 | 4 | 1 | 1 |
| **Type B (N=16)** | - | 1 | 5 | 10 |
| **Type C (N=10)** | - | - | 8 | 2 |

This demonstrates that the two locally-determined ecosystem services act as a stabilizing and simplifying force on the gradient profiles. Their strong signals tend to create clear, linear trends. Removing them unmasks the subtler, more complex patterns of the connectivity-dependent ecosystem services, which are more sensitive to the fine-scale structure of the border. This finding reinforces the necessity of our integrated, multi-service bundle approach, as an analysis based solely on one type of ecosystem service would provide an incomplete picture. For a visual illustration of the on-the-ground landscape patterns associated with the synergies and trade-offs discussed in the main text, see Figure S11.

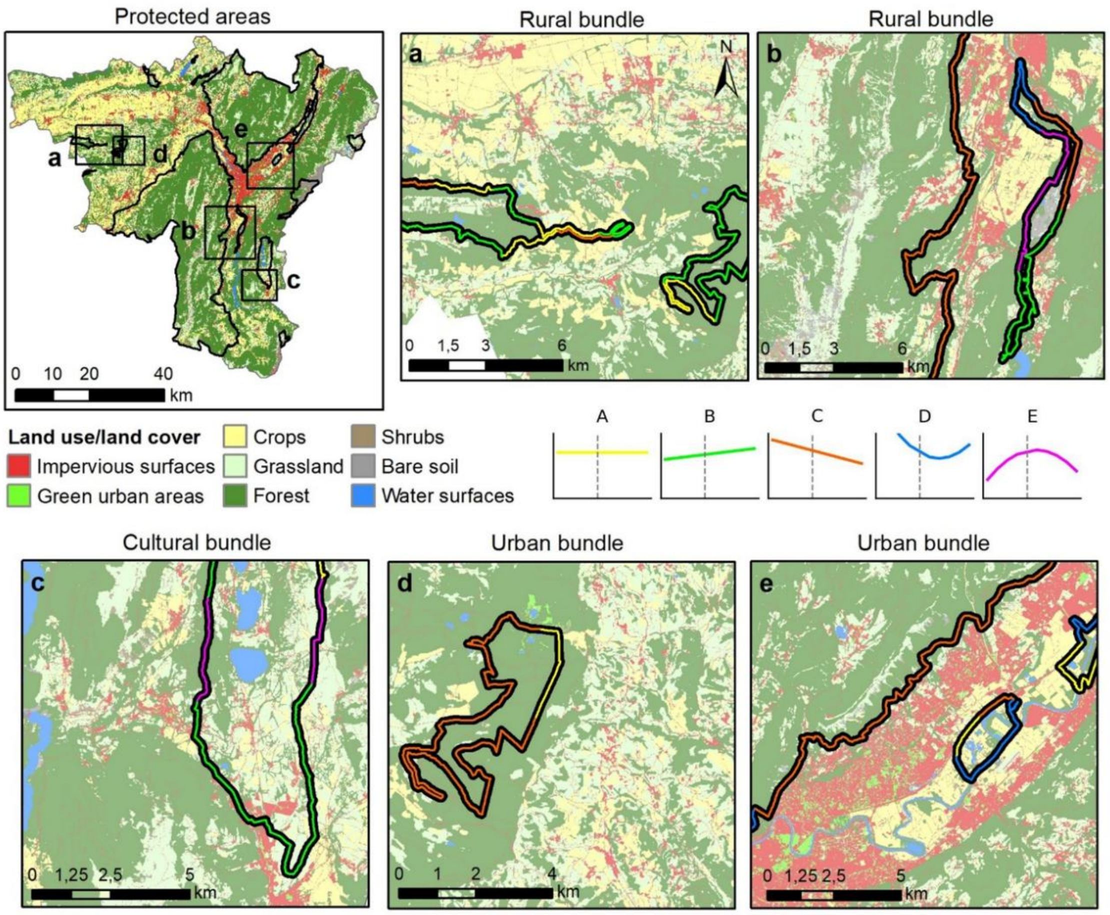


**Figure S 11.** Representative examples of border configurations across varying landscape contexts within the Grenoble region. (a) Rural Bundle: Small protected areas embedded in agricultural landscapes, showing 'Increasing Gradient' (Type B) profiles combined with 'Flat Gradient' (Type A) profiles around two small protected areas. (b) Rural Bundle: Large vs. small protected areas in varied landscapes, contrasting a large protected area border adjacent to an urban area (predominantly 'Decreasing Gradient', Type C) with a smaller protected area border near agriculture in a more heterogeneous landscape (showing 'Increasing Gradients' (Type B), 'Decreasing Gradients' (Type C), 'Boundary Depressions' (Type D) and 'Interface Peaks' (Type E)). (c) Cultural Bundle: A medium-sized protected area in a less intensive landscape, with predominantly 'Increasing Gradient' (Type B) and 'Interface Peak' (Type E) borders. (d) Urban Bundle: A small protected area at the urban-agricultural interface, showing an example of a 'Flat Gradient' (Type A) in the vicinity of a rural mosaic, and a predominance of a 'Decreasing Gradient' (Type C) on the more natural and semi-natural sides. (e) Urban Bundle: Protected areas at the edge of a dense urban area, showing a predominant 'Decreasing Gradient' (Type C) combined with a smaller protected area exhibiting a diversity of border types ('Flat Gradient' (Type A), 'Decreasing Gradient' (Type C), and 'Boundary Depression' (Type D)) within an agricultural and nature corridor.

#### S8. Methodological interpretations and considerations

Our analysis of ecosystem service gradients is focused on characterizing the patterns at the immediate interface between protected areas and their surrounding landscapes. The use of 8 km transects (extending 4 km inwards and outwards) was designed to capture these fine-scale border effects. We therefore acknowledge that the patterns observed, particularly the continuously 'Increasing Gradients' (Type B), are a function of this analytical scale; such a trend would inevitably decline at broader distances. Our study does not model these longer-distance decay curves, but rather provides a high-resolution characterization of the critical zone where management interventions are often concentrated. However, our open-source code is fully adaptable, allowing future studies to modify the transect length to suit different spatial contexts and research questions.

A key innovation of our framework is the integration of spatially-explicit human demand. This approach is most robust for ecosystem services whose beneficiaries are local or regional. We recognize that for ecosystem services like food production and carbon storage, this is a simplification. The demand for these ecosystem services is largely global and delocalized, driven by complex socio-economic dynamics ("telecoupling") that create off-stage burdens and benefits far from the point of provision (Pascual et al., 2017). However, their inclusion is essential. Food production fundamentally shapes the agricultural matrix, while carbon sequestration is not only critical in the context of global climate change but is also the target of numerous international and regional conservation programs. Both ecosystem services, therefore, profoundly influence the local provision of and trade-offs with all other ecosystem services, making them an indispensable component of a realistic border analysis.

Finally, we emphasize that the composition of our three management priorities (rural, urban, and cultural) is specific to the socio-ecological context of the French Alps, informed by prior regional stakeholder consultations (Vannier et al., 2019). This grouping reflects the management priorities and dominant land-use conflicts of this particular region. In other landscapes with different socio-economic drivers or cultural values, the same ecosystem services might be grouped differently, or other ecosystem services might become more relevant. For example, in a region dominated by coastal tourism or forestry, the composition of the cultural and rural bundles would likely change significantly. Our framework is designed to be adaptable, but the interpretation of the results is always contingent on this local context.

We also acknowledge the inherent limitations of the individual ecosystem service models. For instance, our carbon stock model relies on data extrapolated from similar Alpine regions, and the heatwave and flood regulation models were adapted from more urban-focused contexts. While our use of broader biophysical indicators mitigates these limitations, all modeling decisions are documented in the detailed methods sections (Appendix S3), and we recommend that future applications adapt models to their specific contexts.

Several aspects of our border sampling methodology also warrant consideration. The 8 km transect length was designed to capture near-border gradients and may be insufficient for larger study areas. While our adaptive line truncation rules ensure robustness, highly irregular or serpentine-shaped protected areas can present geometric challenges. Furthermore, our method results in a lower transect density at sharp polygon vertices compared to straighter segments. Although we believe this has minimal impact on the overall results, future refinements could incorporate adaptive transect densification at vertices. Despite these potential areas for refinement, we are confident that our current sampling approach effectively captures the ecosystem service variance for the purposes of this study.

Finally, our border analysis is sensitive to the methodological choices in both sampling and classification. The segmentation of borders into units of analysis is a key step, and alternative approaches—such as using different segment lengths, aggregating transects based on underlying landscape characteristics, or aligning segments with administrative boundaries—could yield different insights. Similarly, the classification is inherently sensitive to the selection of the algorithmic thresholds, namely the Mean Squared Error and the flatness threshold. This sensitivity, however, also provides methodological flexibility, allowing these values to be adapted to different management objectives. For instance, we employed a relatively strict flatness threshold. This ensures high confidence in the classification of "Flat Gradients," but it may also mean that some borders with very shallow slopes are classified as Type B or C. Future work could address these sensitivities by testing alternative segmentation strategies or by incorporating slope intensity as a sub-classification criterion, providing even more nuanced information for spatial planning.
